# Comparative metabolomics of fruits and leaves in a hyperdiverse lineage suggests fruits are a key incubator of phytochemical diversification

**DOI:** 10.1101/2021.01.28.427500

**Authors:** Gerald F. Schneider, Diego Salazar, Sherry B. Hildreth, Richard F. Helm, Susan R. Whitehead

## Abstract

Interactions between plants and leaf herbivores have long been implicated as the major driver of plant secondary metabolite diversity. However, other plant-animal interactions, such as those between fruits and frugivores, may also be involved in phytochemical diversification. Using 12 species of *Piper*, we conducted untargeted metabolomics and molecular networking with extracts of fruits and leaves. We evaluated organ-specific secondary metabolite composition and compared multiple dimensions of phytochemical diversity across organs, including richness, structural complexity, and variability across samples at multiple scales within and across species. Plant organ identity significantly influenced secondary metabolite composition, both independent of and in interaction with species identity. Leaves and fruit shared a majority of compounds, but fruits contained more unique compounds and had higher total estimated chemical richness. While organ-level chemical richness and structural complexity varied substantially across species, fruit diversity exceeded leaf diversity in more species than the reverse. Furthermore, the variance in chemical composition across samples was higher for fruits than leaves. By documenting a broad pattern of high phytochemical diversity in fruits relative to leaves, this study lays groundwork for incorporating fruit into a comprehensive and integrative understanding of the ecological and evolutionary factors shaping secondary metabolite composition at the whole-plant level.

## Introduction

Phytochemistry plays a key role in mediating the ecological and evolutionary dynamics of plant interactions (Kessler & Baldwin, 2002; Wittstock & Gershenzon, 2002; Hartmann, 2007). As functional traits, secondary metabolites can significantly affect plant fitness by defending plants against antagonists, directly affecting the competitive ability of neighboring plants, protecting plants from harsh environmental conditions, and attracting and rewarding mutualists, both above and below ground (Iason *et al*., 2012). However, research on secondary metabolites and their role in the ecology and evolution of plants has been disproportionately focused on vegetative organs, specifically the leaf (e.g. Kursar *et al*., 2009; Richards *et al*., 2015; Volf *et al*., 2018; Salazar *et al*., 2018). While secondary metabolites have numerous demonstrated functions mediating plant-animal interactions surrounding leaves, they also likely perform a crucial and complex set of functions in reproductive organs.

Plant reproductive organs have been a nexus of plant-animal interactions since before the emergence of angiosperms. However, the ecological role that secondary metabolites play in the biology of these plant organs has not been deeply explored. Fruits, and the seeds they contain, provide a direct link to plant fitness and are therefore likely to be under intense selection pressure to attract mutualists and deter antagonists. These complex and contrasting selective pressures are distinct from those acting on leaves, and may lead to the occurrence of secondary metabolites not found in other organs. Indeed, given the complex and often contrasting nature of selective pressures to which fruits and seeds are exposed, fruits and seeds are likely to serve as evolutionary incubators of novel secondary metabolites, and disproportionately contribute to the diversity of phytochemical traits. This is especially likely in systems involving animal-mediated seed dispersal (zoochory), in which plants face the ecological and physiological challenge of attracting and offering a nutritional reward to dispersal vectors while also repelling seed predators, pathogens, and non-target frugivores (Herrera, 1982; Tewksbury, 2002; Whitehead *et al*., 2016).

Secondary metabolites endemic to fruits, and with demonstrated functional significance in seed dispersal and/or fruit defense, have been shown in several systems, including iridoid glycosides in honeysuckles (Whitehead & Bowers, 2013a, 2013b), capsaicinoids in *Capsicum* (Suzuki & Iwai, 1984; Tewksbury & Nabhan, 2001; Tewksbury *et al*., 2008), and amides and alkenylphenols in *Piper* (Whitehead *et al*., 2013, 2016; Whitehead & Bowers, 2014; Maynard *et al*., 2020). Further, capsaicinoids in *Capsicum* and alkenylphenols in *Piper* are synthesized only in the fruits of these taxa (Suzuki & Iwai, 1984; Maynard *et al*., 2020). Overall, these studies suggest that unique and potentially contrasting selective pressures on fruits may be an important factor shaping phytochemical diversification in plants. However, our understanding of the relative importance of interactions across plant organs in shaping phytochemical diversity is limited by a paucity of studies that compare chemical composition and metabolomic diversity across plant organs in an ecological context.

Comparative metabolomic studies across plant organs have the potential to greatly expand our understanding of secondary metabolite function and evolution. Given that metabolites may be organ-specific, the location in which they are expressed in the plant (and consequently, the ecological interactions in which they are involved) can provide valuable insight into both the evolutionary origins and ecological consequences of the vast diversity of undescribed plant secondary metabolites.

Despite the likelihood of distinct selective pressures promoting divergent evolution of secondary metabolites across plant organs, it is likely that the phytochemical diversity in one organ may be constrained by physiological or genetic linkages with the phytochemistry of other organs (Adler *et al*., 2006, 2012; Kessler & Halitschke, 2009; Keith & Mitchell-Olds, 2019). Physiological constraints may result when a majority of the steps in a secondary metabolite pathway are localized to a particular part of the plant, yielding complete or nearly complete end products that are then transported to the organs in which they are utilized, e.g. glucosinolates in the Brassicaceae (Keith & Mitchell-Olds, 2019). Such a pathway has a limited capacity to generate organ-specific modifications of its end products prior to transport, and the sink organs may lack the metabolic machinery required for such modifications. Other secondary metabolites are locally synthesized, but in this case organ-specific metabolites derived from a shared metabolic pathway may be limited by genetic linkage, through co-localization of genes responsible for modifications within a metabolic pathway, e.g. terpene synthase clusters (Falara *et al*., 2011; Chen *et al*., 2020; Xu *et al*., 2020). Certainly, evolutionary processes may overcome these constraints when there are conflicting selection pressures among organs, as evidenced by the examples above of compounds occurring only in specific organs. Furthermore, even when fruits and leaves do share compounds, these compounds may be quantitatively uncorrelated (Cipollini *et al*., 2004; Whitehead & Bowers, 2013; Berardi *et al*., 2016). Thus, while all plant species are biochemically circumscribed to some extent by the biosynthetic pathways acquired through their evolutionary history, broad evolutionary patterns of such constraints across plant organs have yet to be elucidated. Comparative metabolomics provide us with the tools to define and characterize these patterns of constraint in conjunction with patterns of phytochemical innovation.

In this study, we use comparative untargeted metabolomics to explore whether and how differential selective pressures and constraints across reproductive and vegetative organs have shaped the diversity and distribution of secondary metabolites in *Piper*, a pantropical species-rich genus. *Piper* are diverse and dominant members of neotropical lowland forest understories and are known to contain a rich array of secondary metabolites (Kato & Furlan, 2007; Richards *et al*., 2015). Their well-studied chemical composition and a long history of ecological research have made them a model system for understanding phytochemical diversification and its role in shaping plant interactions and community structure (Dyer & Palmer, 2004; Richards *et al*., 2015; Salazar *et al*., 2016).

Our overall objective in this study is to test the hypothesis that fruits can act as incubators of phytochemical diversification in plants. First, we describe the occurrence patterns of secondary metabolites across leaves, fruit pulp, and seeds in 12 *Piper* species, providing baseline data for understanding *Piper* secondary metabolite function. We use untargeted mass spectrometry-based metabolomics, molecular networking, and in-silico fragmentation modeling to characterize undescribed metabolites, followed by machine learning and distance-based methods to compare composition across organs and species. Second, we use these data to test predictions of high relative diversity in fruits derived from our hypothesis of fruit-driven phytochemical diversification. We compare multiple dimensions of phytochemical diversity across leaves and fruit organs, including the richness at multiple scales (alpha and gamma diversity), variability (beta diversity), and structural complexity of secondary metabolites.

## Materials and Methods

### Study system

Encompassing over 1,000 species across the Neotropics (Quijano-Abril *et al*., 2006), the genus *Piper* is diverse and abundant in forest understories, clearings, and edges (Gentry, 1990; Dyer & Palmer, 2004). *Piper* growth forms range from herbs and vines to shrubs and small trees (Gentry, 1990; Dyer & Palmer, 2004). Fruits of Neotropical *Piper* are borne on distinct spike-shaped infructescences that are dispersed primarily by bats of the genus *Carollia* (Phyllostomidae). Fruit antagonists of *Piper* include insect seed predators, which have been found to consume up to 87% of seeds (Greig, 1993), and a largely uncharacterized suite of pathogens, which rapidly attack fruit upon ripening (Thies & Kalko, 2004; Whitehead & Bowers, 2014; Maynard *et al*., 2020). Leaves of *Piper* are subject to herbivory from a broad array of arthropods, including a genus of specialist geometrid moths, *Eois*, estimated to include over 1,000 species in the Neotropics (Brehm *et al*., 2016), as well as other geometrid moths, coleopterans, and orthopterans (Dyer & Palmer, 2004).

### Field collections

All field collections took place between 2009 and 2012 at La Selva Biological Station, Heredia Province, Costa Rica. Samples were collected during a phenology census across 28 species of *Piper* during 2009-10 and opportunistically from 2010-12 when ripe fruits were available. Ripe fruits were distinguished by a distinct softening and swelling of the fruit along an infructescence combined with a partial senescence of the infructescence from the branch (presumably to allow bats to easily remove the entire infructescence in flight). In most *Piper* species included in this study, one or a few infructescences ripen per day per plant during the fruiting period, and the vast majority of these are removed on the same night of ripening by bats (Thies & Kalko, 2004; Maynard *et al*., 2020). Those that are not removed rapidly decompose; therefore, we always took care to collect freshly-ripened infructescences. We chose 12 species for inclusion in this study for which we were able to obtain collections from at least three individual plants. For each individual, we collected 1-2 ripe infructescences and the unripe infructescences that were immediately distal to the ripe ones on the same branch. Fruits on a *Piper* branch mature sequentially from the proximal to the distal end of the branch; thus, these adjacent unripe infructescences were the next closest to maturity on that branch. Leaves were collected from the same branch. We chose the youngest fully-expanded leaf that did not have extensive herbivore damage. All samples were transported immediately to the laboratory (within 2 hours) and frozen at -80^°^C prior to analysis. Subsequent analyses involve four sample types: complete leaves, pulp from unripe and ripe infructescences, and seeds from ripe infructescences.

### Chemical extractions

The frozen plant material was freeze-dried (−20° C/ -55 ° C, shelf/condenser), then ground to a fine powder using a FastPrep-24 homogenizer. Seeds and pericarp were separated prior to grinding by gently rubbing the dried fruit over fine mesh; the lignified central rachis of the infructescence was discarded. In unripe fruit, seeds that were not sufficiently developed to be separated from the pericarp by this method were homogenized with the pericarp. For each sample, 50 mg of homogenized powder was weighed into a 2 mL Eppendorf tube using a microbalance. To isolate the broadest possible range of phytochemicals while excluding the broadest possible range of primary metabolites, extracts were prepared using buffered acetonitrile and acetone in series. The acetonitrile and acetone extraction solutions were prepared with an aqueous acetate buffer (44.3 mmol/L ammonium acetate), both at 70:30 solvent: buffer, v/v. The solutions were prepared with Nanopure® water, Fisher HPLC-grade acetic acid, and Fisher Optima®-grade ammonium acetate, acetonitrile, and acetone. All containers and instruments coming into contact with the extracts were rinsed with Fisher Optima®-grade methanol. Each 50 mg sample was extracted twice with 1.5 mL buffered acetonitrile, then twice more with 1.5 mL buffered acetone (6.0 mL total extraction solution). During each of these four extractions, the sample was mixed with the extraction solvent for 5 min in a vortexer, and then centrifuged for 5 min at 15870 rcf, after which the supernatant was removed and added to a 20 mL glass scintillation vial. The supernatant from each of the four extractions was combined in the same 20 mL vial. The combined extract was dried at 30° C using a nitrogen evaporator until no solvent was visible, then further dried in a lyophilizer for 12 h (−20° C/ -55 ° C, shelf/condenser) before being transferred to storage at -80° C until analysis.

### Untargeted metabolomics

LC-MS data were collected using an Acquity I-class UPLC coupled to a Waters Synapt G2-S quadrupole time-of-flight mass spectrometer (Waters). For analysis, dried extracts were resuspended at 10 mg/mL in 75:25 water: acetonitrile + 0.1 % formic acid, with 1.0 µg/mL *N*-oleoylglycine as an internal standard. The extract was then sonicated for 10 min, after which a 20 µL aliquot was taken and diluted 10-fold with 75:25 water: acetonitrile + 0.1 % formic acid. The diluted aliquot was then vortexed and centrifuged (10 min,13,000 xg) and an aliquot (180 µL) was transferred to an LC-MS vial for analysis. Solvent blanks and combined, quality-control samples were injected at regular intervals during data collection. The autosampler temperature was 10°C and the injection volume was 1.5 µL. The column employed was a reverse-phase Acquity BEH C18 (2.1 mm ID x 150 mm, 1.7 um particle size, Waters) maintained at 35 °C at a flow rate of 0.2 mL/min. Solvent A was water with 0.1% formic acid and solvent B was acetonitrile with 0.1% formic acid (LCMS grade, Fisher Chemical). Solvent gradient: 0-0.5 min, 90% A; 0.5-1.0 min, 75% A; 1.0-8.0 min, 5% A; 8.0-10.0 min, held at 5% A; 10.0-11.0 min, 90% A; 11.0-15.0 min, held at 90% A. Mass spectra and fragmentation spectra were collected simultaneously using Waters’ MS^E^ in positive-ion mode, with the following parameters: peak data recorded in centroid mode; 0.185 s MS scan time; 20-35 V collision energy ramp; argon collision gas; 125° C source temperature; 3 V capillary voltage; 30 V sample cone voltage; 350° C desolvation temperature; nitrogen desolvation at 500 L/hr; 10 µL/min lockspray flow rate; 0.1 s lockspray scan time; 20 s lockspray scan frequency; 3 lockspray scans to average; 0.5 Da lockspray mass window; 3 V lockspray capillary voltage. The lockspray solution was 1 ng/ µL leucine enkephalin, and sodium formate was used to calibrate the mass spectrometer.

Alignment, deconvolution, and annotation of molecular and adduct ions were conducted using the XCMS and CAMERA packages in R statistical software (Smith *et al*., 2006; Tautenhahn *et al*., 2008; Benton *et al*., 2010; Kuhl *et al*., 2012).

### Molecular networking

Molecular networking was used to quantify and visualize the dimensions of the chemical structural trait space occupied by the secondary metabolites in our study (Aron *et al*., 2020). This technique employs tandem mass spectrometry to generate fragmentation spectra for each putative compound. These fragmentation spectra are diagnostic of molecular structure, and through pairwise comparison they are used to generate a network linking putative compounds to one another based on structural similarity.

In our study, fragmentation spectra data files were aligned, deconvoluted, and converted to .mgf using MS-DIAL software (v4.10) and were then uploaded to the Global Natural Products Social Molecular Networking (GNPS) online workflow for molecular networking and library-based annotation. The following parameters were used for the GNPS workflow METABOLOMICS-SNETS-V2 (v14): 0.02 Da precursor ion mass tolerance; 0.02 Da fragment ion mass tolerance; minimum matched fragment peaks = 6; minimum cluster size = 3; minimum cosine score for network pairs = 0.7; network TopK = 1000; maximum connected component size = 0. All mass spectral libraries available through GNPS which contained data collected in positive ion mode were used for annotation. Library search parameters were: minimum matched peaks = 6; cosine score threshold = 0.6; maximum analog mass difference = 100. Workflow options for advanced filtering, advanced GNPS repository search, and advanced output were not used.

For further annotation via *in-silico* modeling, results of the METABOLOMICS-SNETS-V2 workflow were passed to a second GNPS workflow, Network Annotation Propagation (NAP_CCMS v1.2.5). The parameters used for NAP_CCMS were as follows: all clusters selected; subselection cosine value = 0.7; first candidates for consensus score = 10; fusion results used for consensus; accuracy for exact mass candidate search = 15 ppm; acquisition mode = positive; adduct ion types = [M+H]^+^ and [M+Na]^+^; all structure databases selected; no custom database or parameter file; compound class not specified; parent mass selection enabled; maximum number of graphed candidate structures = 10; standard workflow type.

Finally, the outputs from METABOLOMICS-SNETS-V2 and NAP_CCMS were combined and exported for visualization using the GNPS workflow MolNetEnhancer (v15). Network visualization and curation was conducted using Cytoscape software (v3.7.2). Parent masses of features in the molecular network were curated based on the XCMS-CAMERA output described above, with primary metabolites and artefactual or pseudoreplicated features removed from the network and subsequent analyses. Features in the molecular network were annotated to the level of chemical class, e.g. flavonoid or prenol lipid, based on ClassyFire chemical taxonomy as applied by MolNetEnhancer. The list of annotated molecular features returned by XCMS-CAMERA processing was used to compare overall phytochemical composition across organs and species.

Unfragmented ions collected during single-mass-spectrometry and subsequently aligned, deconvoluted, and annotated, as described above, were used to compare overall phytochemical composition across organs and species. Ion abundance data were transformed to presence/absence data using the peak recognition parameters in XCMS (R code repository). Ion presence/absence was used for analyses rather than relative ion abundance for two reasons: 1) our sample size affords limited capacity to account for variation in abundance within a given organ of a given species, and 2) the scale of variation in ion abundance is likely to differ widely across the structurally diverse compounds in *Piper* due to variation in ionization efficiency (Cech & Enke, 2001).

### Comparisons of phytochemical composition across organs and species

To compare metabolome-level patterns of phytochemical composition across organs and species, we conducted two separate analyses of the multivariate sample composition, focused first on compound occurrences (presence/absence data) and second on the structural composition of samples. First, to visualize differences in patterns of compound occurrence across samples, we used non-metric multidimensional scaling (NMDS) based on the Sørensen dissimilarity index (binary Bray-Curtis). We then tested for effects of organ, species, and their interaction on compound composition using PERMANOVA, implemented with the ‘adonis2’ function in the R package ‘vegan’. The individual plant identity was included in these analyses as a ‘strata’ (i.e. random effect), and we used 999 permutations (note that this means the minimum possible *P*-value is *P* = 0.001, indicating that the observed differences in sample composition could not be replicated in any of the 999 permutations). To further understand specific differences among the four organ types, we followed this analysis with post-hoc pairwise PERMANOVAs for all possible combinations of organ types, correcting for multiple comparisons using the ‘pairwise.adonis2’ function (Martinez Arbizu, 2020). In addition, based on strongly supported interactions between organ and species (see results), we also divided the data by species and tested for the effects of organ on compound composition for each species individually. All analyses were conducted using the ‘vegan’ package in R (Oksanen *et al*., 2019).

In addition to our analysis of compound occurrence, we also examined how the structural composition of samples was affected by organ, species, and their interaction. To account for structural features, we generated a multivariate structural dissimilarity index that was a modification of Sedio *et al*.’s (2017) Chemical Structural and Compositional Similarity (CSCS) index, which quantifies the pairwise similarity of samples by calculating the maximum cosine similarity of the aligned MS-MS ion fragmentation spectra for each inter-sample pair of molecular features. We modified this index by representing ion abundance as a binary term and expressing the index in terms of dissimilarity (1-CSCS). The structural dissimilarity matrix was then used as the basis for NMDS and PERMANOVAs as above that examined the effects of organ, species, and their interaction on structural composition.

### Machine learning

To identify molecular features that distinguished different organs, we used random forest analysis via the “randomForest” and “Boruta” packages for R statistical software (Liaw & Wiener, 2002; Kursa & Rudnicki, 2010). All molecular features distinguished in XCMS-CAMERA processing were used as variables in these analyses. The random forest analysis used a decision tree model to assign samples to our four organ groups (Breiman, 2001; 2002). In the process, the analysis ranked molecular feature variables according to their importance in the model’s group assignments. Boruta analysis complemented the random forest analysis by applying a search for molecular features that were important in informing group assignments. This is accomplished by comparing the features’ importance with importance achievable at random, using “shadow” variables which are generated by permuting the original variables (Kursa & Rudnicki, 2010).

### Comparisons of chemical diversity across organs

Phytochemical diversity is a multifarious concept that includes the number of compounds (richness), their relative abundances (evenness), their structural complexity, and their variation in space and time (Wetzel & Whitehead, 2020). Considering the challenges associated with estimating abundances in untargeted LC-MS-MS data, we focus here on richness and structural complexity, both of which were examined at multiple scales within and across species. For each organ type, we define gamma diversity as the total diversity observed across all samples for that organ, alpha diversity as the average diversity within a single sample from one organ from one *Piper* individual, and beta-diversity as the variation (both intra- and inter-specific) across samples.

#### Gamma diversity

To compare the gamma diversity (total number of compounds detected across all species) of different organs, we used a rarefaction analysis analogous to those commonly used to assess species diversity (Gotelli & Colwell, 2011) with compounds as “species” as in Wetzel & Whitehead (2020). This allowed us to: 1) explicitly visualize the relationship between chemical diversity and sampling scale across different organs (i.e. alpha, beta, and gamma diversity), and 2) estimate the total compound richness in each organ type. Because our individual samples were not independent (we collected three samples per species for 12 species), we used a constrained rarefaction that is similar conceptually to spatially-constrained rarefaction (Chiarucci *et al*., 2009). Briefly, samples were added to bootstrapped accumulation curves in a semi-random manner in which samples from the same species were grouped. For each iteration, a random sample was chosen as a starting point, then other samples from that species were added in random order prior to choosing another sample at random, following with all other samples from that species, and so on until all species were included. We estimated total species richness from these curves using the ‘fitspecaccum’ function in ‘vegan’ based on an asymptotic regression model. Accumulation curves and fits were averaged across 5000 bootstrapped samples with random starting points.

#### Alpha diversity

To compare the average compound richness in a sample (i.e. alpha diversity) across organs, we used a linear mixed model with organ, species, and their interaction as fixed effects and plant identity as a random effect. For hypothesis testing, we compared the full model to simplified versions with fixed effects terms deleted using likelihood ratio tests. Based on a strong interaction between organ and species (see results), we further divided the data by species and examined differences in richness among organs for each species separately.

#### Structural complexity

To compare structural complexity across organ types, we first calculated an index of structural complexity for each sample that was modified from the CSCS index described in Sedio *et al*. (2017) to include only presence/absence data. This within-sample CSCS represents the mean pairwise similarity among all individual molecular features detected in a sample. We used the inverse of this similarity index (1-CSCS) as a measure of overall structural complexity present in a sample. To examine how structural complexity varied across organs and species, we used a linear mixed model with species, organ, and their interaction as fixed effects and plant identity as a random effect. Hypothesis testing was conducted as described above using likelihood ratio tests. Based on strong interactions between organ and species (see results), we examined differences among organs separately for each *Piper* species.

#### Beta diversity

We examined differences in beta-diversity (i.e. sample-to-sample variance in composition) across organs in two ways, focusing first on variation in compound occurrences (presence/absence) and second on structural features. These analyses were based on the same distance matrices described above that we used to assess overall differences in composition across samples, but instead focused on variance (i.e. dispersion) among samples. This was assessed using the function ‘betadisper’ in the R package ‘vegan’ to compare the dispersion around the group centroid across the four organ types. The ‘betadisper’ function calculated the distances from each sample to the group centroid, and statistical support for differences in dispersion across organs was assessed using a permutation test (N = 999 permutations) followed by a post-hoc Tukey HSD test to assess pairwise differences among individual organs. Because this analysis focused on sample-to-sample variance and our dataset included multi-level sampling (multiple species and multiple individuals within species), significant differences in beta diversity across organs could be due to both intraspecific and interspecific variance among samples. Thus, we followed this analysis with a set of PERMANOVAs, conducted separately for each organ type, with *Piper* species as an explanatory factor. This analysis allowed us to test if *Piper* species explained a significant portion of the variation in composition within an organ, and partitioned sample-to-sample variance within an organ type according to the percent of variance explained by species and the percent explained by differences among individuals within species (i.e. the residual variance).

## Results

### Untargeted metabolomics and molecular networking reveal high chemical diversity and many compounds unique to fruits

Alignment, deconvolution, and annotation of molecular and adduct ions via XCMS and CAMERA yielded 1,311 unique molecular features across all species and organs. Most of these compounds (1,126) occurred in all organ types, 92 of these were unique to fruits (unripe pulp, ripe pulp, and/or seeds) and 4 were unique to leaves (Fig. 1a). It is important to note that, like all other metabolomic approaches, our analytical approach is likely to overestimate the true number of individual chemical compounds present in our samples. The combination of XCMS-CAMERA followed by manual curation unfortunately cannot condense all features (m/z and retention time pairs) into individual compounds. In-source fragmentation, ion clusters, centroid peak splitting of highly abundant ions, and centroid merging of ions near the noise level can all contribute to expanding the dataset beyond individual compounds. The 1,311 features described in this work thus overestimates the number of individual molecular species, though to a lesser extent than in uncurated datasets. Nevertheless, this overestimation is likely to represent a small fraction of the total chemical diversity captured in our analysis. Furthermore, this overestimation is also likely to be of equal magnitude across all species and organs and therefore, will not have a significant impact on the general conclusions of our study. Regarding terminology, these 1,311 features meet or exceed the level of curation beyond which features have, for clarity, been described as “compounds” in the chemical ecology literature (e.g.: Sedio *et al*., 2017; Christian *et al*., 2020; Ricigliano *et al*., 2020). Thus, for the sake of consistency and clarity, we refer to our curated features as compounds.

**Figure 1:**
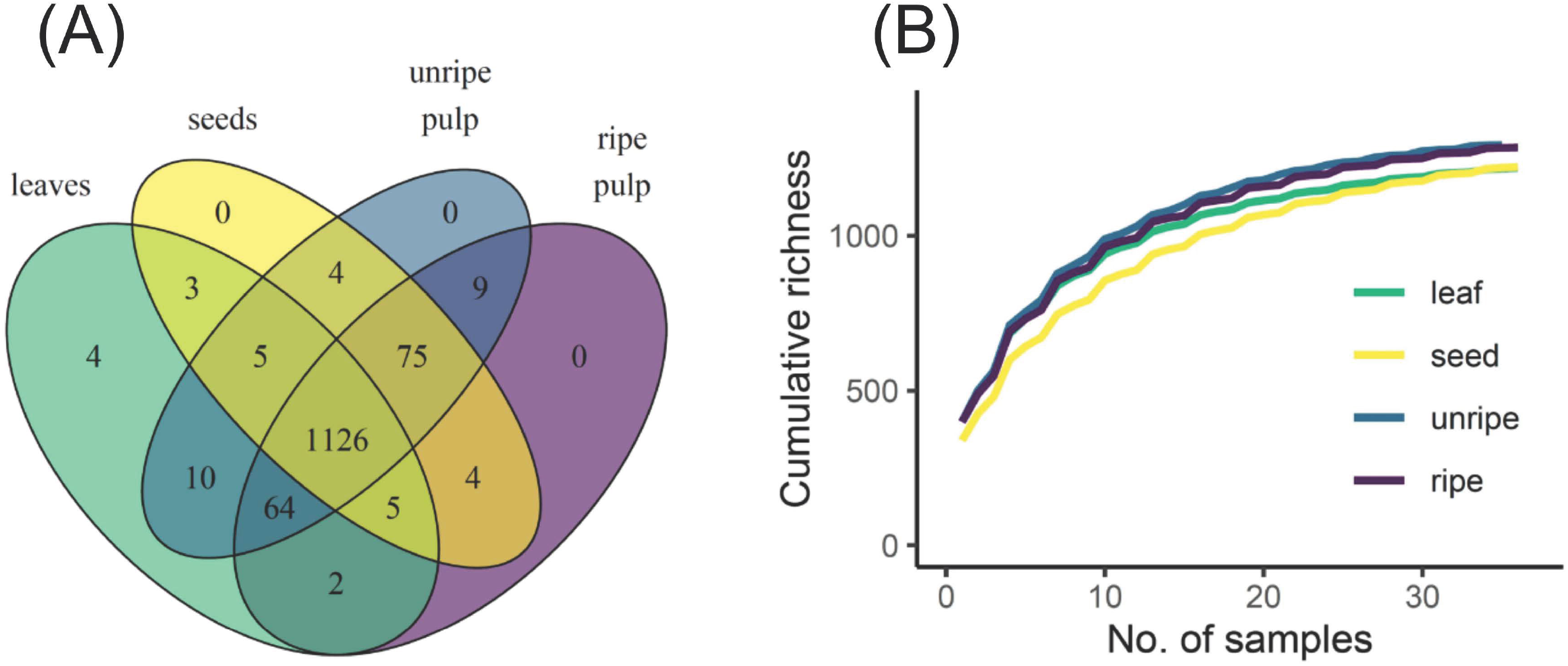
Chemical gamma diversity parsed by organ type. A Venn diagram (**A**) shows the total number of compounds detected across all samples that were unique and shared across organ type. The rarefaction curve (**B**) shows how compound richness accumulates with sampling scale in each organ type. Curves represent an average across 5000 bootstrapped accumulation curves with random starting points. Because samples from the same species were not independent, the rarefaction was constrained by species such that samples from the same species were always added in sequence.

Tandem mass spectrometry yielded fragmentation spectra for 706 of these compounds (Table 1, Fig. 2). Library- and *in silico*-based classification of fragmentation spectra and parent ions via GNPS resulted in annotation at the level of “class” *sensu* ClassyFire chemical taxonomy for 527 compounds in 23 classes and 179 were unknowns (Table 1, Fig. 2). Of the 706 compounds that yielded fragmentation spectra, 62 were unique to fruits and 2 were unique to leaves (Table 1). Fruit-specific richness was higher than leaf-specific richness across all chemical classes, these differences were statistically supported for carboxylic acids and derivatives and prenol lipids (Table 1; Fig. 2).

**Table 1:** Summary of GNPS molecular network annotations. Compound richness indicates the number of putative compounds, for which fragmentation spectra were obtained, that fall under the given category. Chemical classes are per ClassyFire chemical taxonomy. Chemical classes which represented ≤ 1% of all annotated compounds were categorized as “Other”. A compound was labeled as fruit or leaf-specific if it was detected only in that organ within the 12 focal *Piper* species. Asterisks indicate statistical support (P < 0.05) for differences between fruit-specific and leaf-specific richness (binomial test with probability = 0.5).

| Chemical Class | Examples of class known from *Piper* spp. | Total Compound Richness | Fruit-specific Compound Richness | Leaf-specific Compound Richness |
| --- | --- | --- | --- | --- |
| Benzene and substituted derivatives | Cyanogenic benzoates, Non-prenylated benzoic acids | 75 | 5 | 0 |
| Carboxylic acids and derivatives | Amides, Chromenes, Kavalactones | 122 | 25* | 1 |
| Flavonoids | Flavonoids | 104 | 3 | 0 |
| Organo-oxygen compounds | Oxygenated or glycosidic derivatives of other classes | 65 | 5 | 0 |
| Other | Amides, Chalcones, Chromenes, Imides | 37 | 4 | 0 |
| Prenol lipids | Chalcones, Prenylated benzoic acids, Neryl<br>catechol diols,<br>Terpenes | 124 | 6* | 0 |
| Unknown |  | 179 | 14* | 1 |
| Total |  | 706 | 62* | 2 |

**Figure 2:**
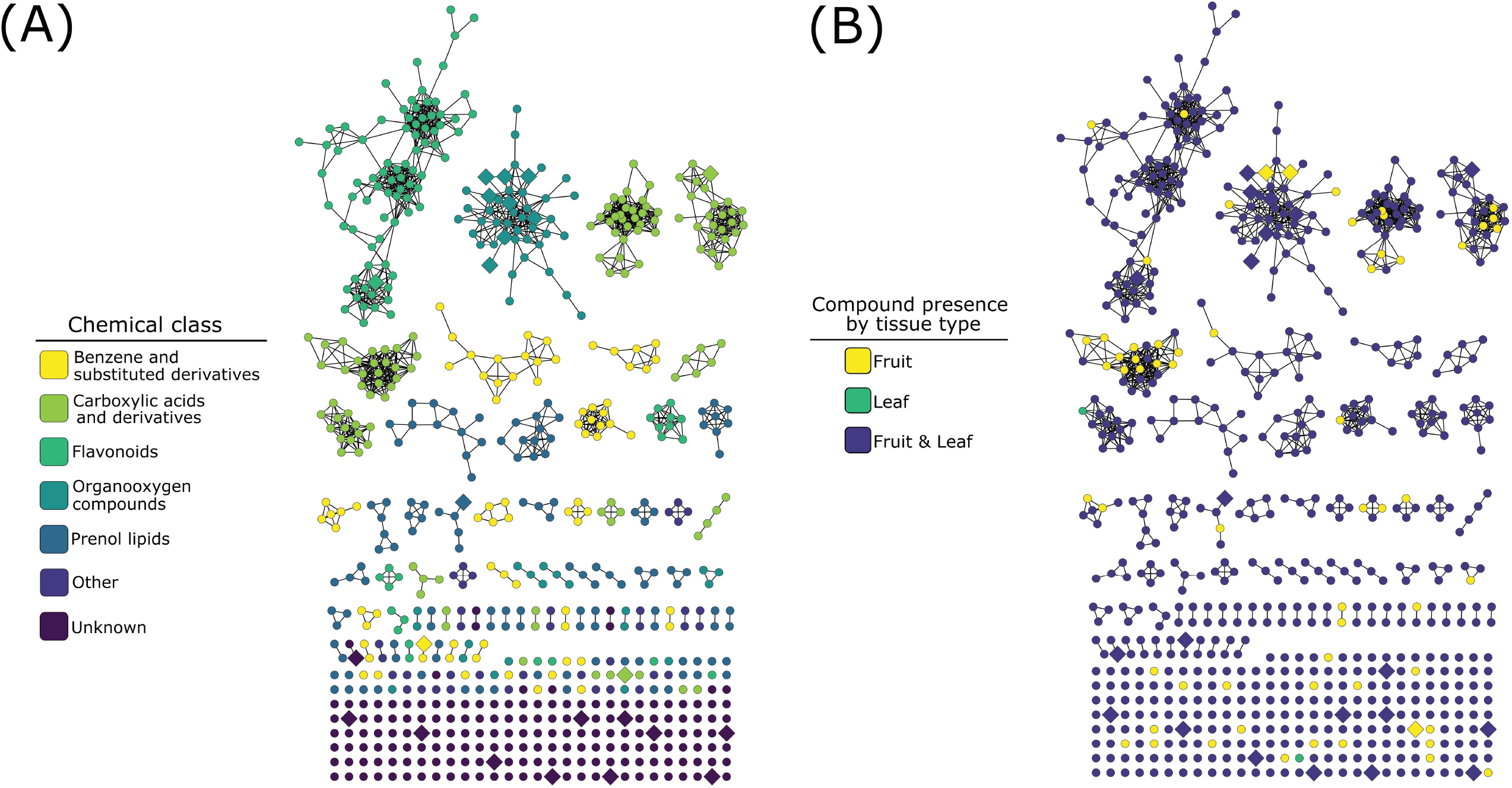
Molecular network of 706 compounds from 12 *Piper* species color-coded by ClassyFire chemical classification annotation (**A**) or by organ-level occurrence across the 12 species (**B**). Node and edge arrangement and compound annotation are as described in “Molecular Networking” methods. Enlarged, diamond-shaped nodes represent compounds identified by the Boruta analysis as important for distinguishing among organs. In (**B**), compounds are coded as occurring in “fruit” if they occur in one or more of the three sample types (unripe pulp, ripe pulp, or seeds).

### Phytochemical composition differs across organs and species

The multivariate patterns of phytochemical occurrence were strongly affected by organ, species, and their interaction (organ: *F*_*3,95*_ = 24.19, *P* = 0.001; species: *F*_*11,95*_ = 27.65, *P* = 0.001; organ x species: *F*_*33,95*_ = 1.99, *P* = 0.001; Fig. **3a**). Pairwise comparisons among organs indicated strong differences among all organs (*P* = 0.001 for all comparisons). Further examination of differences among organs for each of the 12 *Piper* species individually also revealed strong effects of organ in all cases (Table 2). Similarly, when we assessed factors influencing the multivariate patterns of structural composition across samples, we found a strong effect of organ, species, and their interaction (organ: *F*_*3,95*_ = 17.34, *P* = 0.001; species: *F*_*11,96*_ = 21.28, *P* = 0.001; organ x species: *F*_*33,96*_ = 2.31, *P* = 0.001; Fig. **3b**), significant differences among organs in all pairwise comparisons (*P* = 0.001 for all comparisons), and differences among organs for each individual species (Table 2).

**Table 2:** Results from PERMANOVAs, conducted separately for each species, testing the effects of organ type (leaves, seed, unripe pulp, or ripe pulp) on two aspects of phytochemical composition: compound occurrences and structural composition

| *Piper* species | Compound Occurrence |  | Structural Composition |  |
| --- | --- | --- | --- | --- |
| | $F_{3,11}$ | $P$ | $F_{3,11}$ | $P$ |
| aduncum | 3.07 | 0.001 | 3.07 | 0.001 |
| auritum | 3.51 | 0.002 | 3.51 | 0.002 |
| biolleyi | 2.19 | 0.009 | 2.19 | 0.009 |
| colonense | 4.39 | 0.001 | 4.39 | 0.001 |
| generalense | 4.11 | 0.001 | 4.11 | 0.002 |
| glabrescens | 4.18 | 0.002 | 4.18 | 0.001 |
| multiplinervum | 3.45 | 0.001 | 3.45 | 0.001 |
| peltatum | 4.50 | 0.001 | 4.50 | 0.001 |
| reticulatum | 5.38 | 0.001 | 5.38 | 0.002 |
| sancti-felicis | 3.67 | 0.001 | 3.67 | 0.001 |
| silvivagum | 5.19 | 0.001 | 5.19 | 0.001 |
| umbricola | 2.89 | 0.002 | 2.89 | 0.001 |

### Machine learning, informed by numerous compounds from diverse chemical classes in each organ, accurately distinguishes between reproductive and vegetative organs

The random forest decision tree model used 2000 trees with 36 variables at each split. Our analysis showed an overall out-of-bag (OOB) mean error rate of 11.72% across the four organ groups. In other words, using secondary metabolites alone, the algorithm was able to predict if a sample was from a leaf, ripe fruit, unripe fruit, or seed approximately 9 times out of every 10 samples. Examining the error rate of each organ group, it was apparent that correctly assigning pulp samples to the correct ripeness stage was the main source of OOB error, with error rates of 27.78% and 18.92% for unripe and ripe pulp respectively. Leaves and ripe seeds both exhibited zero OOB error. Boruta analysis, designed to both identify important classification features and assess their relative contribution to the final classification performance, identified 23 features exhibiting a significantly higher variable importance score (VIS) than shadow variables. These 23 features are detailed in Table S1.

### Comparisons of chemical diversity across organs

#### Gamma diversity

Of the 1,311 compounds detected across all organ types, 1,126 were shared across all organs, 92 were found only in fruit (unripe pulp, ripe pulp, and/or seeds), and four were found only in leaves. Rarefaction analysis indicated that the observed richness of secondary metabolites was approaching the asymptote for all organ types (Fig. 1b). Further, this analysis showed that the estimated total gamma diversity (total number of compounds across all 12 species of *Piper*) was highest in unripe and ripe fruit pulp, intermediate in seeds, and lowest in leaves, with only unripe and ripe pulp exhibiting overlapping confidence intervals (Fig. 1b, Table 3).

**Table 3:** Rarefaction results showing total estimated richness for each organ across all 12 Piper species sampled

| Organ | Observed Richness | Estimated Richness | SE | 95% CI high | 95% CI low |
| --- | --- | --- | --- | --- | --- |
| leaf | 1219 | 1226 | 1.5 | 1229.3 | 1223.5 |
| seed | 1222 | 1276 | 3.0 | 1282.0 | 1270.1 |
| unripe pulp | 1293 | 1312 | 2.0 | 1315.8 | 1307.9 |
| ripe pulp | 1285 | 1311 | 2.7 | 1316.5 | 1306.1 |

#### Alpha diversity

In our analysis of average differences in compound richness across organs and species, we found a strong interaction between organ and species (*X*^*2*^ = 128.99, *P* < 0.0001) and further examined differences among organs for each species separately. Organs often showed clear differences in average richness, but the patterns were highly variable across species (Fig. 4). In three species (*P. glabrescens, P. reticulatum*, and *P. slivivagum*), pulp and/or seeds had higher compound richness than leaves. However, in two species (*P. multiplinervum* and *P. generalense*) leaves had higher compound richness than all other fruit organs.

**Figure 3:**
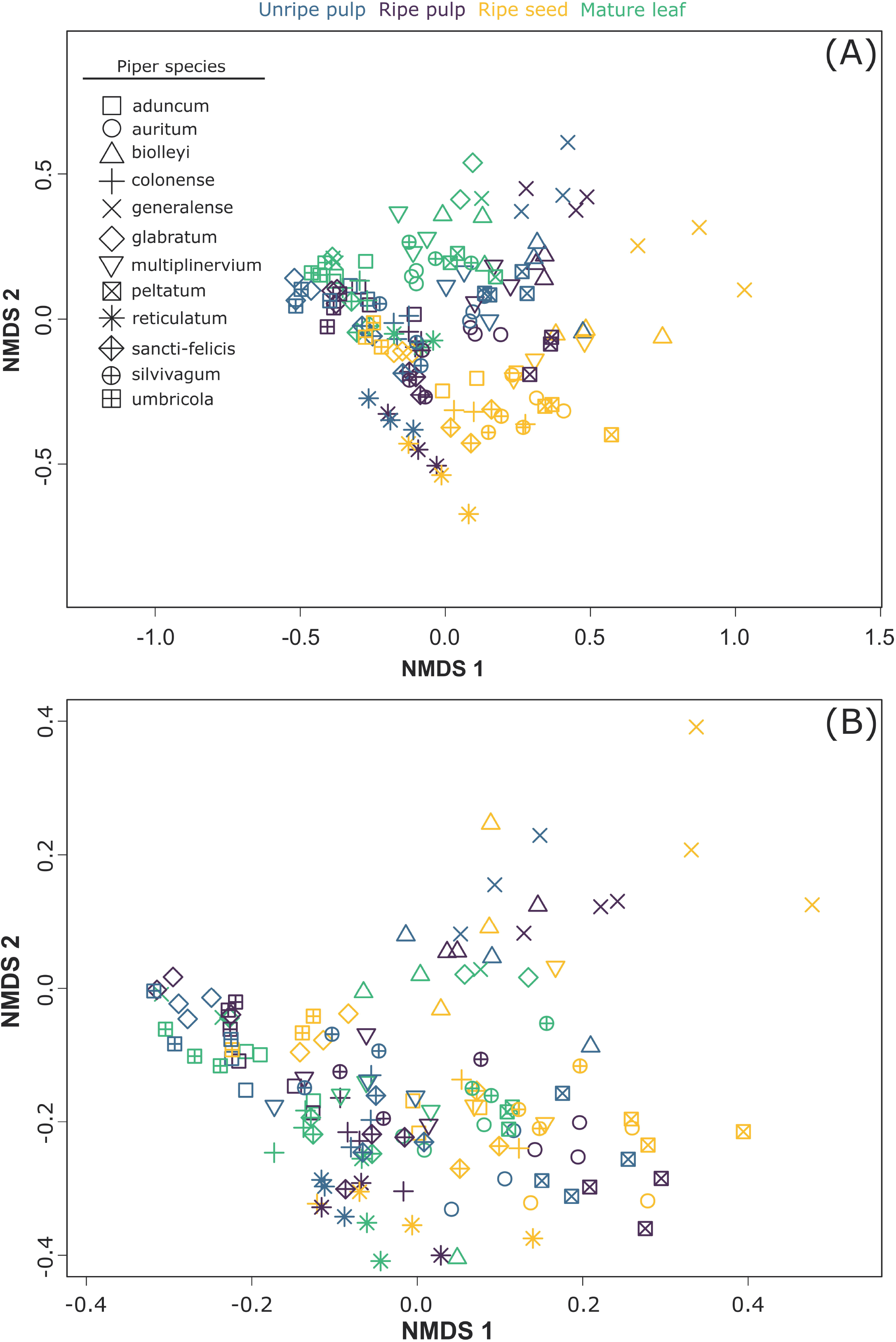
NMDS plots showing the effects of organ and species on two aspects of multivariate chemical composition across samples: (**A**) compound occurrences (presence/absence) and (**B**) structural composition.

**Figure 4:**
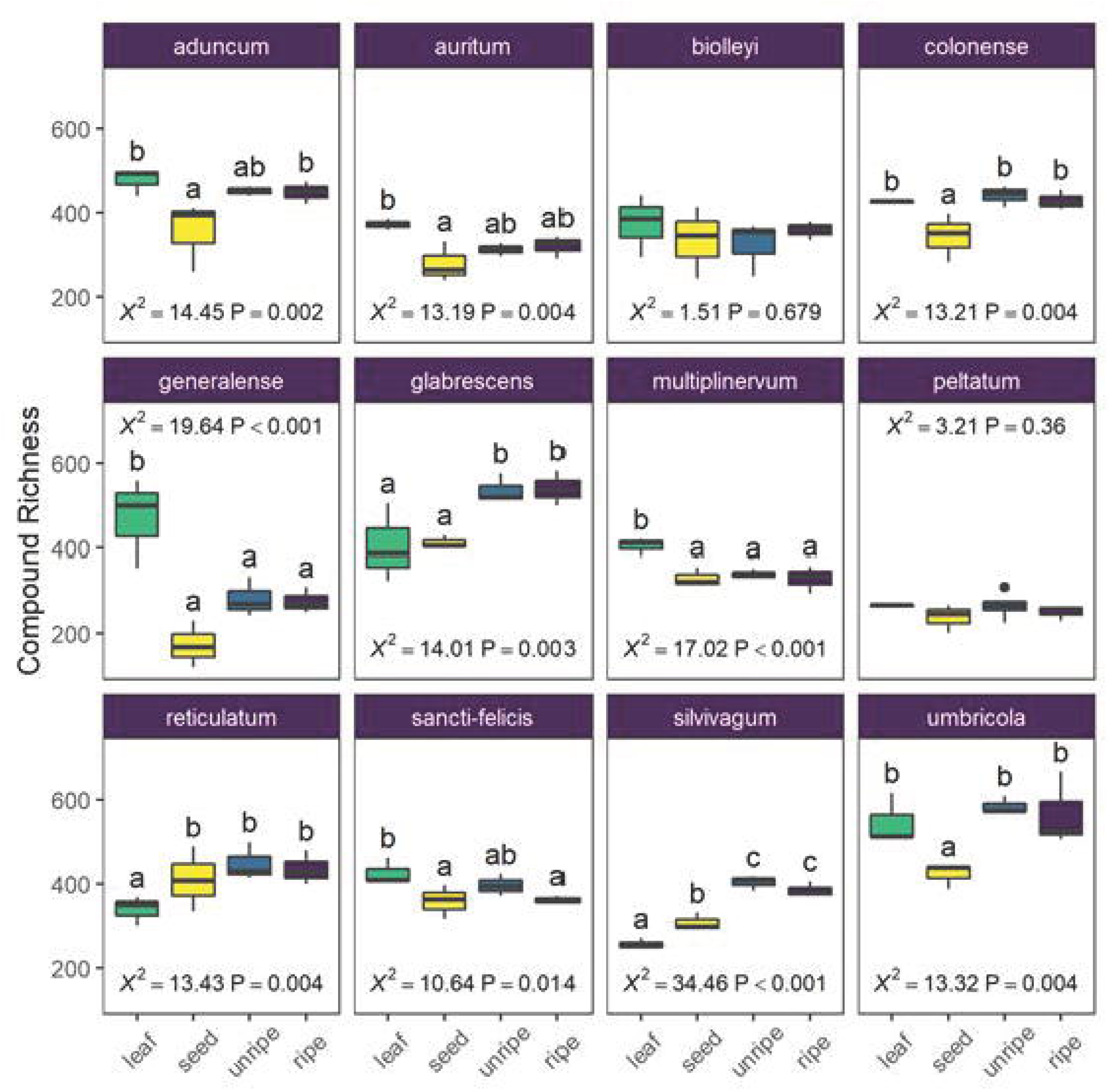
Average chemical richness differs across species and organ type (leaf, seed, unripe pulp, and ripe pulp). Letters indicate results of pairwise Tukey post-hoc comparisons of organs within each species, with non-shared letters indicating a significant difference at *P* < 0.05. Each species plot includes □ ^2^ and *P*-values from species-level LMMs.

#### Structural complexity

In our analysis of average differences in structural complexity across organs and species, we found a strong interaction between organ and species (*X*^*2*^ = 131.13, *P* < 0.0001) and further examined differences among organs for each species separately. For seven of twelve species, organs showed differences in average structural complexity, but the patterns were variable across species (Fig. 5). In two species (*P. glabrescens* and *P. slivivagum*), one or more fruit organs had higher complexity than leaves. In another species (*P. generalense*), leaves had higher complexity than all other fruit organs. Often, seeds had the lowest structural complexity, or at least lower structural complexity than unripe or ripe fruit pulp (Fig. 5).

**Figure 5:**
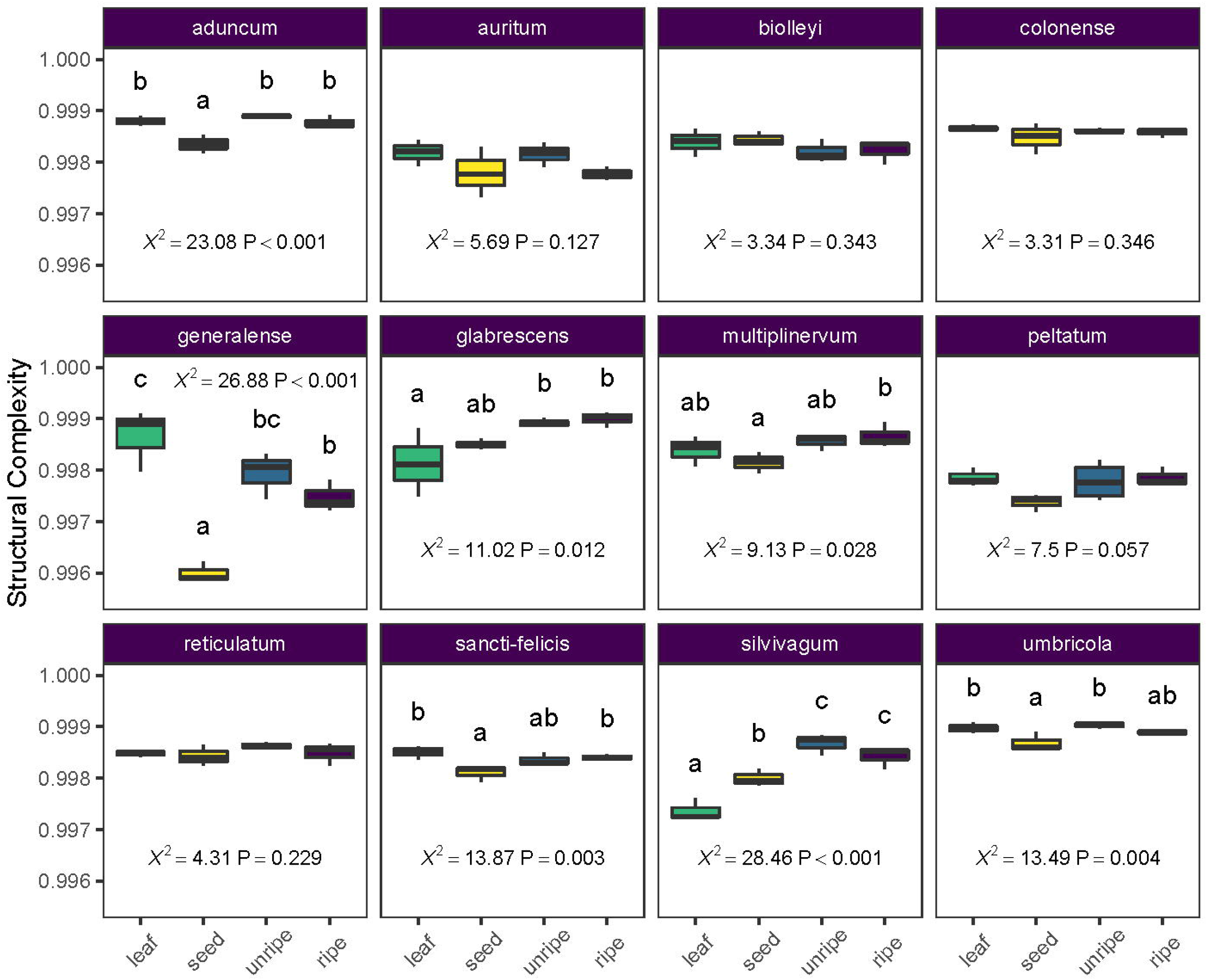
Average structural complexity differs across species and organ type (leaf, seed unripe pulp, and ripe pulp). Letters above each box plot column indicate results of pairwise Tukey post-hoc comparisons of organs within each species, with non-shared letters indicating a significant difference at *P* < 0.05. Each species plot includes □ ^2^ and *P*-values from species-level LMMs.

#### Beta diversity

We found that beta-diversity in chemical composition was higher for fruits than leaves when considering only compound occurrences as well as structural composition. First, for compound occurrences, there was strong support for overall differences in beta diversity across organ types (F_3,139_ = 7.56, *P* = 0.001), with higher average distances to the group centroid for seeds, unripe pulp, and ripe pulp relative to leaves (Fig. **6a**). Next, for structural composition, there was also strong support for overall differences in beta diversity across organ types (F_3,139_ = 4.10, *P* = 0.009). In this case, leaves had lower beta diversity than seeds or ripe pulp, and unripe pulp was intermediate (Fig. **6b**). Further analyses conducted separately for each organ type showed that the differences in beta-diversity among organ types was due to variation both at the interspecific and intraspecific level (Table 4). A large proportion of the sample-to-sample variation within organ types (67-86%) was explained by differences among species relative to that explained by variation within species (14-33%), and this was especially true for unripe and ripe fruit pulp (Table 4).

**Table 4:**
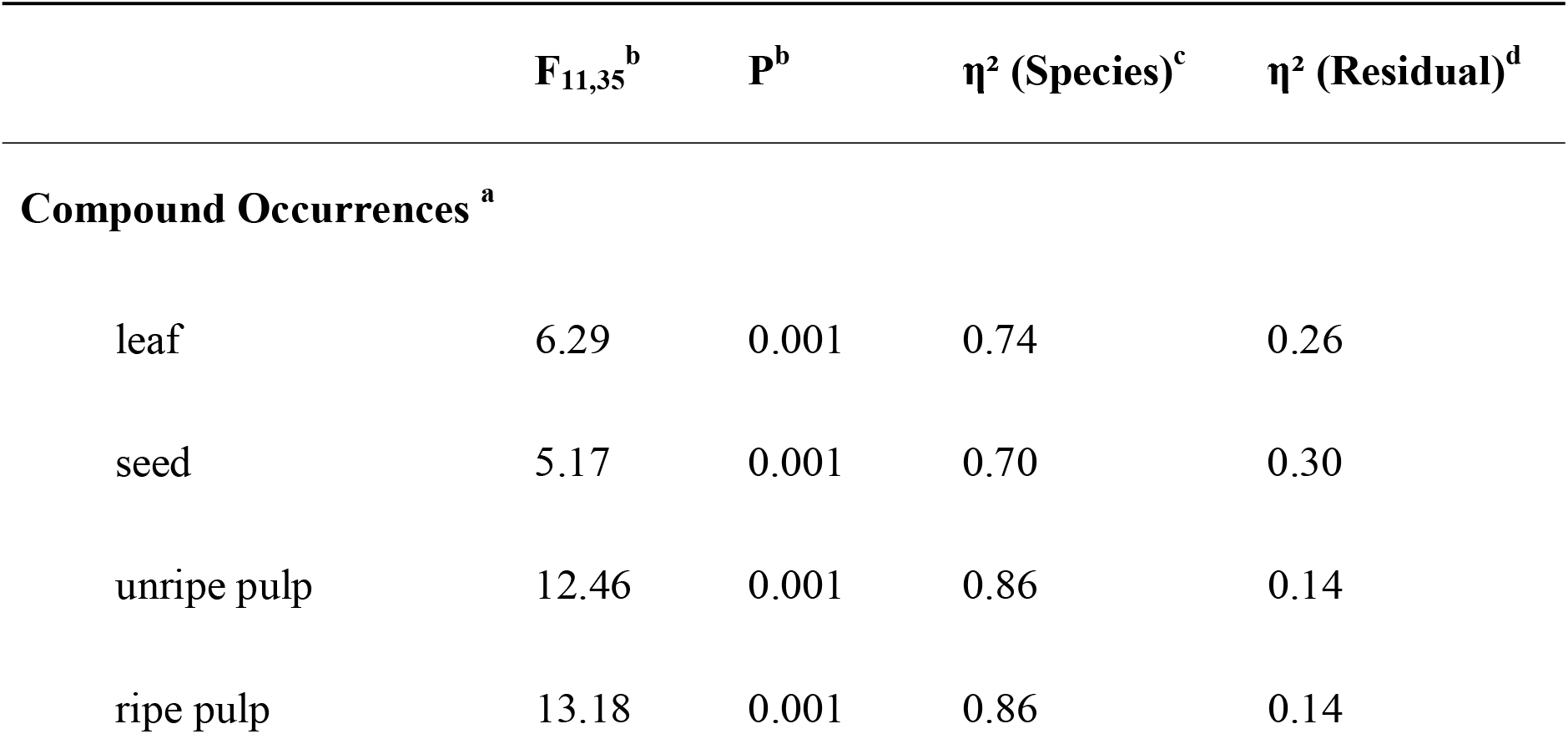
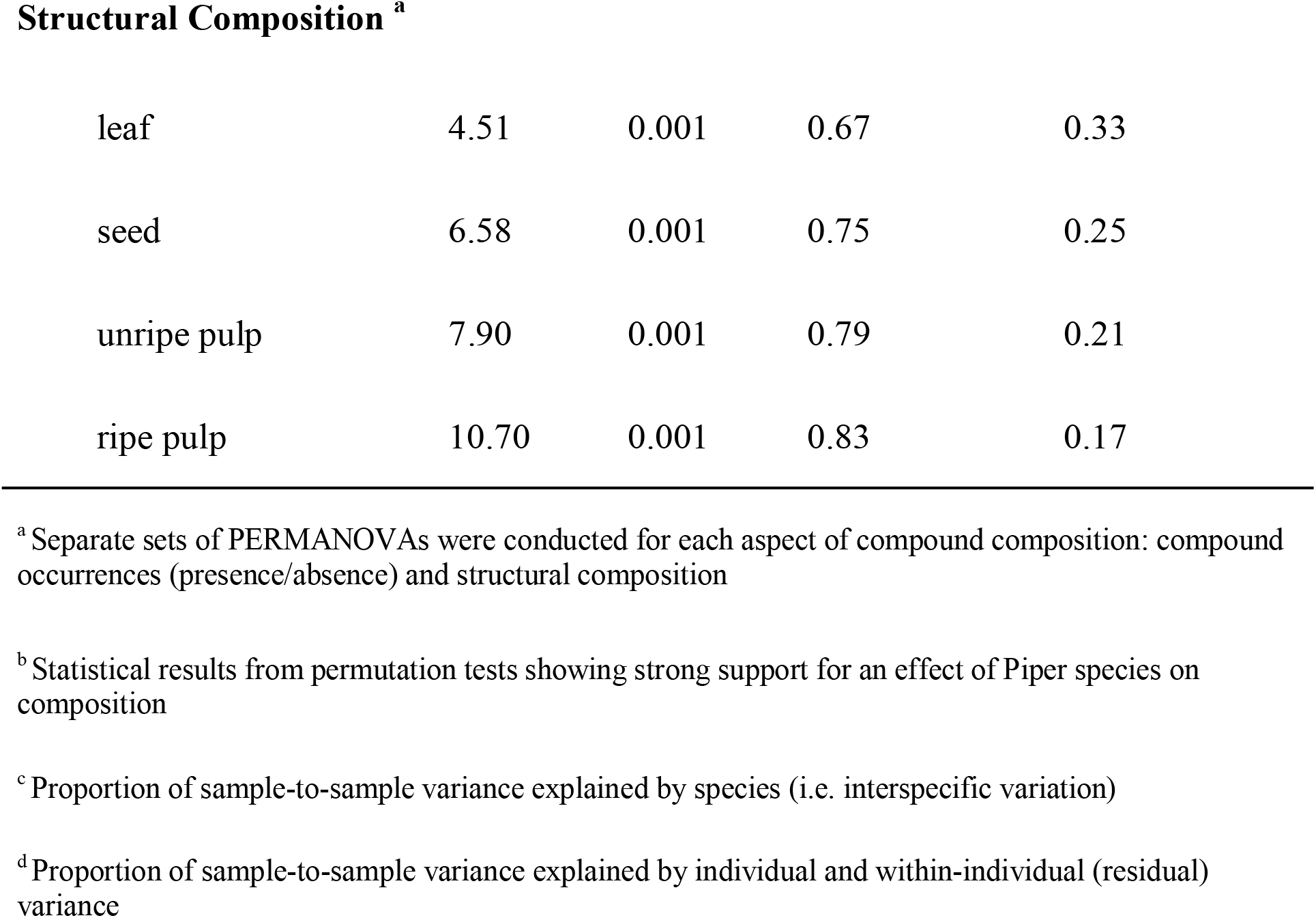
Results from PERMANOVAs showing a large percentage of sample-to-sample variance in composition within organ types (i.e. beta diversity) is explained by species

| | $F_{11,35}^b$ | $P^b$ | $\eta^2$ (Species)<sup>c</sup> | $\eta^2$ (Residual)<sup>d</sup> |
| --- | --- | --- | --- | --- |
| **Compound Occurrences <sup>a</sup>** |  |  |  |  |
| leaf | 6.29 | 0.001 | 0.74 | 0.26 |
| seed | 5.17 | 0.001 | 0.70 | 0.30 |
| unripe pulp | 12.46 | 0.001 | 0.86 | 0.14 |
| ripe pulp | 13.18 | 0.001 | 0.86 | 0.14 |
| **Structural Composition <sup>a</sup>** |  |  |  |  |
| leaf | 4.51 | 0.001 | 0.67 | 0.33 |
| seed | 6.58 | 0.001 | 0.75 | 0.25 |
| unripe pulp | 7.90 | 0.001 | 0.79 | 0.21 |
| ripe pulp | 10.70 | 0.001 | 0.83 | 0.17 |

<sup>a</sup> Separate sets of PERMANOVAs were conducted for each aspect of compound composition: compound occurrences (presence/absence) and structural composition
<sup>b</sup> Statistical results from permutation tests showing strong support for an effect of *Piper* species on composition
<sup>c</sup> Proportion of sample-to-sample variance explained by species (i.e. interspecific variation)
<sup>d</sup> Proportion of sample-to-sample variance explained by individual and within-individual (residual) variance

## Discussion

Secondary metabolites occur in all plant parts, but theory in chemical ecology has largely focused on interactions between leaves and leaf herbivores. In this study, we surveyed and compared the compositional and structural diversity of secondary metabolites across vegetative and reproductive organs in 12 species of the genus *Piper*. Most metabolites were shared across organs, but fruits contained many more unique compounds than leaves. Furthermore, relative to leaves, fruits contained a higher total number of compounds (gamma diversity) and a higher sample-to-sample variance in composition (beta diversity). While *Piper* species varied in terms of whether fruits or leaves had a higher average compound richness per sample (alpha diversity), richness was more often higher in fruits than in leaves. Taken together, these patterns reveal that, in neotropical *Piper*, fruits are overall more chemically diverse than leaves and point to a key role for mutualistic and antagonistic fruit-frugivore interactions in shaping phytochemical evolution and diversity.

Our untargeted metabolomic survey of phytochemical occurrence patterns revealed that fruit organs harbor fruit-specific metabolites from a variety of chemical classes (Table 1, Fig. 2), in each class equal to or greater in number than those that were leaf-specific (Table 1). This included classes of compounds that have previously been found to be more numerous and abundant in fruit organs (e.g. amides; Whitehead *et al*., 2013), as well as numerous chemical classes previously described in studies of *Piper* spp. leaf chemistry (Parmar *et al*., 1997; Baldoqui *et al*., 1999; Kato & Furlan, 2007; Richards *et al*., 2015). The occurrence of numerous fruit-specific secondary metabolites from a variety of unlinked biosynthetic pathways suggests a pattern of fruit-specific secondary metabolite trait evolution, likely a result of fruit-specific selective pressures.

The evolution of organ-specific phytochemical traits across our target plant species is also made evident by the results of our machine learning analysis. Here, our random forest model was very successful at distinguishing among organ types based solely on their secondary metabolite composition. Most notably, the exceptional performance of the classification algorithm to distinguish between vegetative and reproductive organs can only be explained by the presence of strong association between chemical composition and organ type. Despite the fact that our species set included vines, understory shrubs, and pioneering taxa, all adapted to very different local habitats, these associations are consistent across all 12 focal species.

The clustering patterns found by our NMDS analysis (Fig. 3) show clustering at two different levels. First, the samples from different organs from the same species cluster together. This pattern strongly suggests the presence of physiological or genetic linkage constraints in the organ specific evolution of phytochemicals. The strong chemical similarity across organs within a species could point toward the influence that changes in the expression or composition of secondary compounds in one plant organ could have on the expression or composition of other organs. Although our data do not allow us to disentangle the precise mechanisms that give rise to these patterns, it is clear that chemical changes in one plant organ are likely to be mirrored, to some extent, by changes in the chemical architecture of the whole plant. Second, as expected, and despite the strong chemical similarity exhibited by organs within a *Piper* species, samples also show a clear pattern of clustering by organ type (Fig. 3). This pattern reinforces the expectation that the distinctive regimes of selective pressures imposed upon the different plant organs are sufficiently strong to create convergent organ-specific patterns of the chemical composition, and that these selective regimes are likely to be consistent across species and habitats. The Boruta variable importance model, a widely used machine learning algorithm designed to identify statistically important classification variables from large datasets, revealed specific compounds from at least six different chemical classes as key features that distinguish vegetative and reproductive organs (Table S1).

While the majority of significant Boruta variables, like the overall majority of secondary metabolites cataloged in our study, exhibited some overlap in occurrence across leaf and fruit organs when the 12 *Piper* species were evaluated as a group (Fig. 1a), there was substantially less overlap at the level of individual species (Fig. S1). In many cases, these patterns of variance were the result of numerous compounds occurring in only one organ type in a certain species or subset of species, but occurring more widely in another species or subset of species.

The broad overlap across organs in compound occurrence at the genus level provides a degree of insight into the extent of constraints on organ-specific chemical trait evolution at this taxonomic scale. However, to a degree this overlap can also be attributed to the shared demand for defensive compounds across vegetative and reproductive organ types. While phylogenetic data will be required in order to infer the ancestral organ localizations of phytochemical traits of *Piper*, the widespread variation in organ localizations that we observed across species suggests that genetic constraints have not bound these traits to a certain organ type over the course of *Piper* speciation. Further, the apparent mobility of secondary metabolite traits across organ types within the genus suggests a bidirectional exchange of these traits, which, when vegetative and reproductive organs are each threatened by separate assemblages of consumers, may allow more rapid defense trait adaptation than can arise from novel mutations.

Our untargeted metabolomic survey has shown that fruit organs are at the very least a reservoir of phytochemical richness. While the alpha diversity of organs at the species level was highly variable (Fig. 4), rarefaction analysis of gamma diversity showed a small but clear trend towards higher richness of secondary metabolites in reproductive organs (Fig. 1b; Table 3). Similarly, while chemical structural complexity of organs at the species level was highly variable (Fig. 5), chemical structural variance (□ -diversity) across species was significantly higher for reproductive organs than for leaves (Fig. 6). In summary, these trends indicate not only that reproductive organs accumulate a higher number of secondary metabolite traits than do leaves, but also that these traits are more divergent from one another across species than those of leaves. These trends are consistent with higher overall evolutionary diversification of phytochemical traits in reproductive organs, suggesting that fruits may be an important, but underappreciated, force in shaping chemical trait evolution at the whole plant level.

**Figure 6:**
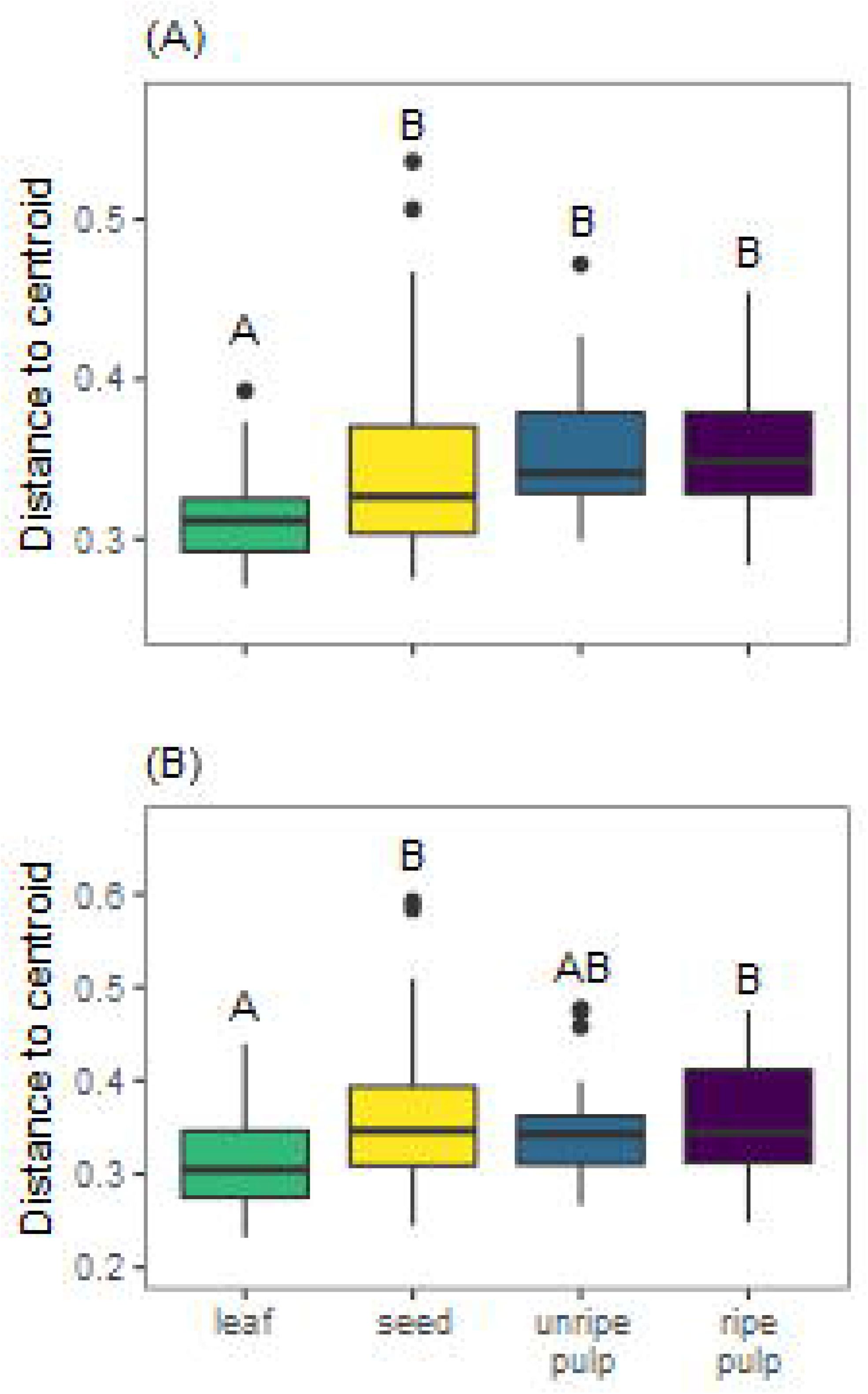
Beta diversity in chemical composition is higher for reproductive organs than leaves when considering variance in compound occurrences (**A**) or structural composition (**B**). Letters indicate results of pairwise Tukey post-hoc comparisons of organs, with non-shared letters indicating a significant difference at *P* < 0.05.

## Acknowledgements

This research was supported by National Science Foundation (Grants No. DEB-1210884 and DEB-1856776 to SRW) and start-up funds to SRW from the Virginia Tech Department of Biological Sciences. The mass spectrometry resources used in this work were maintained with funds from the Fralin Life Science Institute as well as the Virginia Agricultural Experiment Station Hatch Program (VA-160085). We thank the Organization for Tropical Studies for logistical support at La Selva Biological Station. Marisol Luna Martinez assisted with sample collection and Orlando Vargas Ramírez assisted with plant identification. Natalie Rodeman and Katherine Berg assisted with sample processing for chemical analyses.

## Author contributions

GFS, SRW, and DS designed the research; SRW collected field samples; GFS, SBH, and RFH conducted chemical analyses, GFS conducted molecular networking and data curation; GFS and SRW conducted the statistical analysis with contributions from DS; GFS wrote the first draft of the manuscript with contributions from SRW, and all authors contributed substantially to revisions and approved the final version.

## Data availability

UPLC-MS data and associated programming scripts will be made publicly available through Global Natural Products Social Molecular Networking (https://gnps.ucsd.edu/) and all other data and programming scripts that support the findings of this study will be made publicly available at Zenodo https://www.zenodo.org/ upon publication of this manuscript.

