## Supplemental information for "Comparative metabolomics of fruits and leaves in a hyperdiverse lineage suggests fruits are a key incubator of phytochemical diversification"

**Table S1:** Random Forest - Boruta variable importance summary. Chemical classification is per ClassyFire as previously described. Occurrences of compounds with VIS significantly greater than shadow variables were summed for each organ across all 3 samples in all 12 species, i.e. each compound has a maximum occurrence sum of 36 for each organ. As compounds were rarely restricted to only one organ, compounds occurring in more than one organ were counted in each of the organs in which they occurred.

| Chemical Class | m/z ([M+H]$^+$) | Sum of occurrences of compounds with VIS > shadow | | | | Variable importance score |
|---|---|---|---|---|---|---|
| | | Leaf | Unripe pulp | Ripe pulp | Ripe seed | |
| Benzene and substituted derivatives | 362.097 | 22 | 13 | 32 | 29 | 3.6 |
| Carboxylic acids and derivatives | 274.140 | 0 | 1 | 5 | 23 | 2.9 |
| Carboxylic acids and derivatives | 275.124 | 9 | 17 | 32 | 31 | 3.7 |

| | | | | | | |
|---|---|---|---|---|---|---|
| Flavonoids | 311.092 | 22 | 36 | 26 | 22 | 2.3 |
| Organooxygen compounds | 281.071 | 27 | 18 | 32 | 35 | 2.4 |
| Organooxygen compounds | 288.108 | 11 | 14 | 28 | 34 | 3.6 |
| Organooxygen compounds | 319.150 | 2 | 7 | 19 | 29 | 4.8 |
| Organooxygen compounds | 323.145 | 10 | 8 | 20 | 34 | 3.5 |
| Organooxygen compounds | 343.091 | 2 | 13 | 14 | 32 | 4.0 |
| Organooxygen compounds | 466.204 | 5 | 3 | 8 | 28 | 4.6 |
| Organooxygen compounds | 467.188 | 3 | 4 | 10 | 28 | 3.2 |
| Organooxygen compounds | 522.203 | 35 | 21 | 33 | 32 | 3.7 |
| Prenol lipids | 535.271 | 35 | 19 | 21 | 11 | 3.4 |
| Unknown | 220.170 | 34 | 6 | 2 | 0 | 7.4 |
| Unknown | 269.114 | 10 | 8 | 8 | 32 | 3.6 |
| Unknown | 301.144 | 9 | 20 | 6 | 34 | 5.8 |
| Unknown | 318.300 | 0 | 26 | 16 | 36 | 6.4 |
| Unknown | 322.161 | 2 | 1 | 3 | 30 | 5.2 |
| Unknown | 439.142 | 36 | 19 | 24 | 24 | 3.8 |
| Unknown | 568.427 | 33 | 12 | 12 | 5 | 4.7 |
| Unknown | 613.483 | 33 | 12 | 11 | 8 | 5.0 |
| Unknown | 797.518 | 32 | 19 | 17 | 3 | 4.2 |
| Unknown | 954.614 | 25 | 17 | 7 | 1 | 3.0 |
| **Sum** | | **397** | **314** | **386** | **541** | |

**Figure S1:** Molecular network visualizations for individual species. Two network visualizations are shown for each of the 12 *Piper* species analyzed. Network visualizations are shown in alphabetical order, by species. In each pair of visualizations, the above is color-coded by chemical class, while the below is color-coded by organ-level compound occurrence. Node and edge arrangement and compound annotation are as described in "Molecular Networking" methods. Enlarged, diamond-shaped nodes represent compounds identified by the Boruta analysis as important for distinguishing among organs. Compounds are coded as occurring in "fruit" if they occur in one or more of the three sample types (unripe pulp, ripe pulp, or seeds). Compounds not present in a species are shown with high transparency node shading (chemical class visualization) or gray node shading ( organ occurrence visualization).

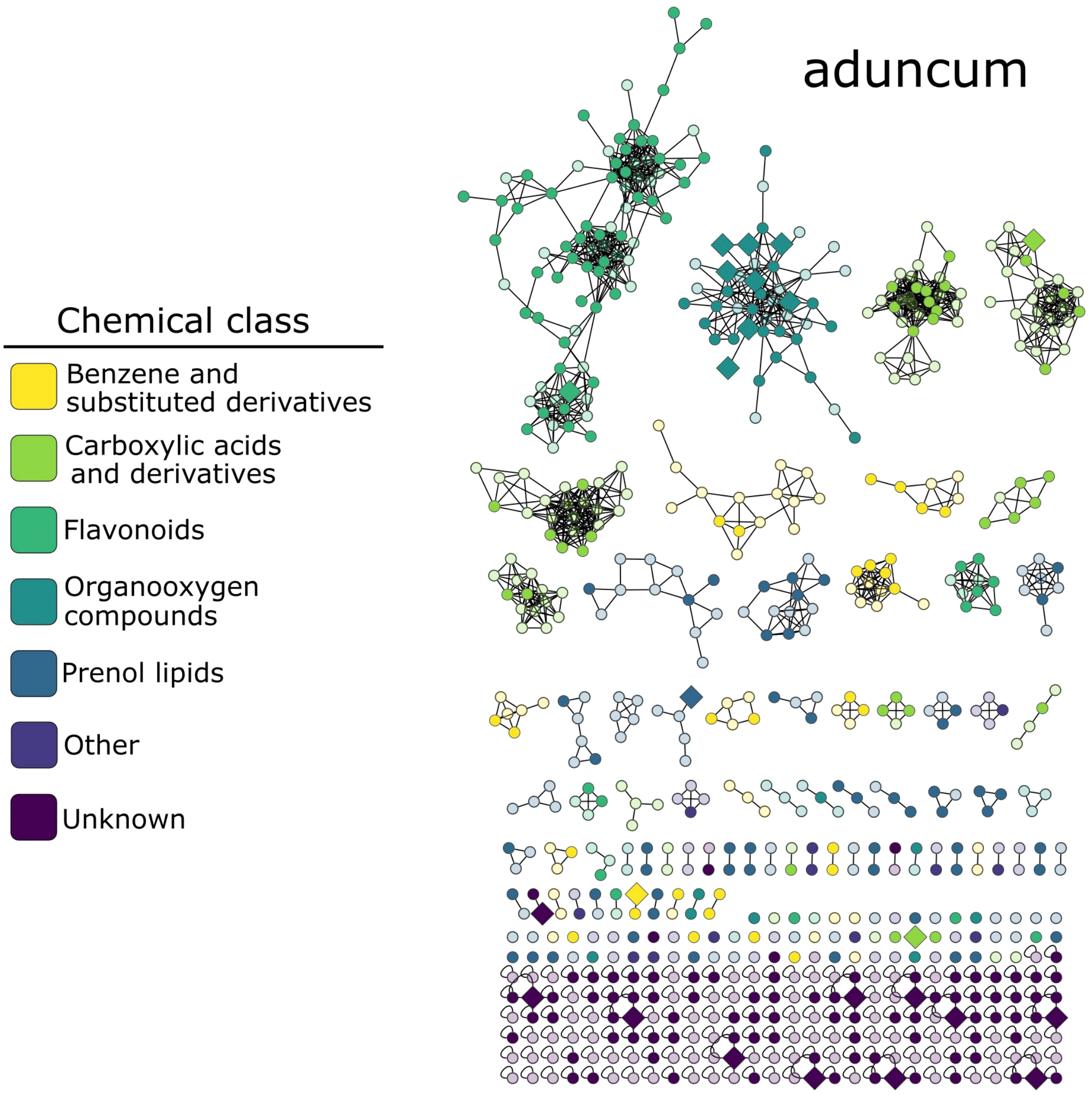

Chemical class
Benzene and substituted derivatives
Carboxylic acids and derivatives
Flavonoids
Organooxygen compounds
Prenol lipids
Other
Unknown
aduncum


29

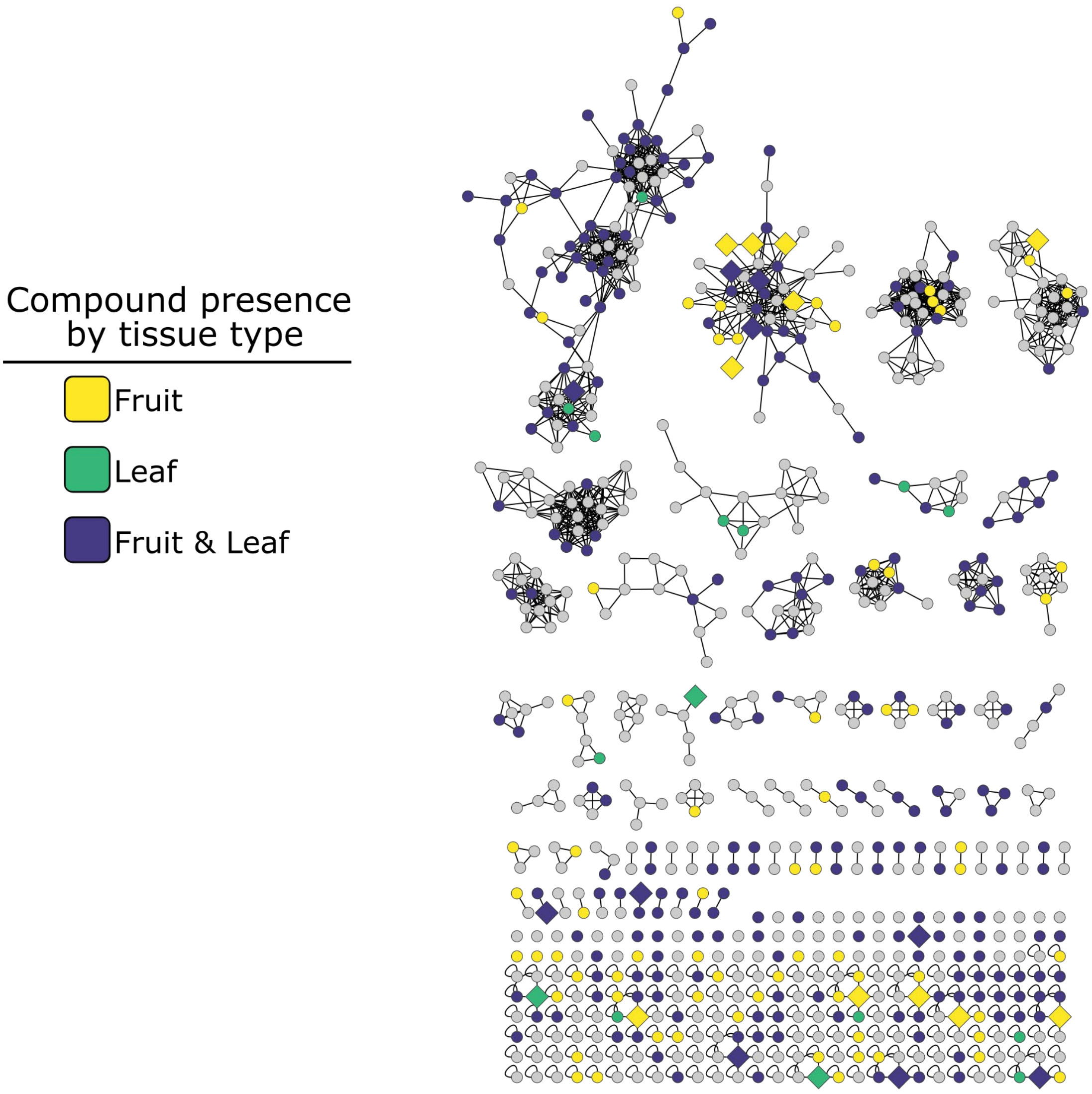
Compound presence
by tissue type
Fruit
Leaf
Fruit & Leaf

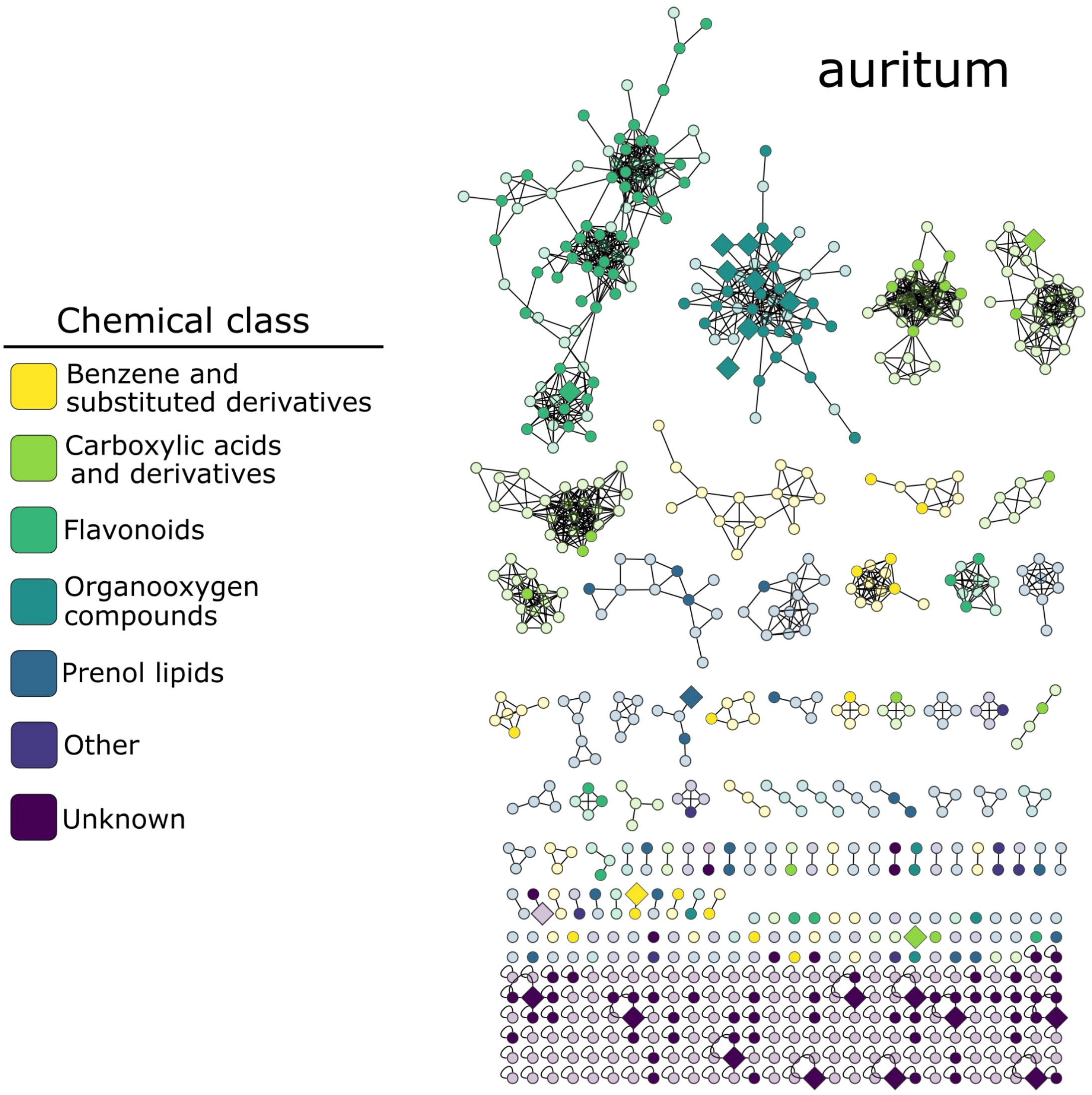
auritum
Chemical class
Benzene and substituted derivatives
Carboxylic acids and derivatives
Flavonoids
Organooxygen compounds
Prenol lipids
Other
Unknown

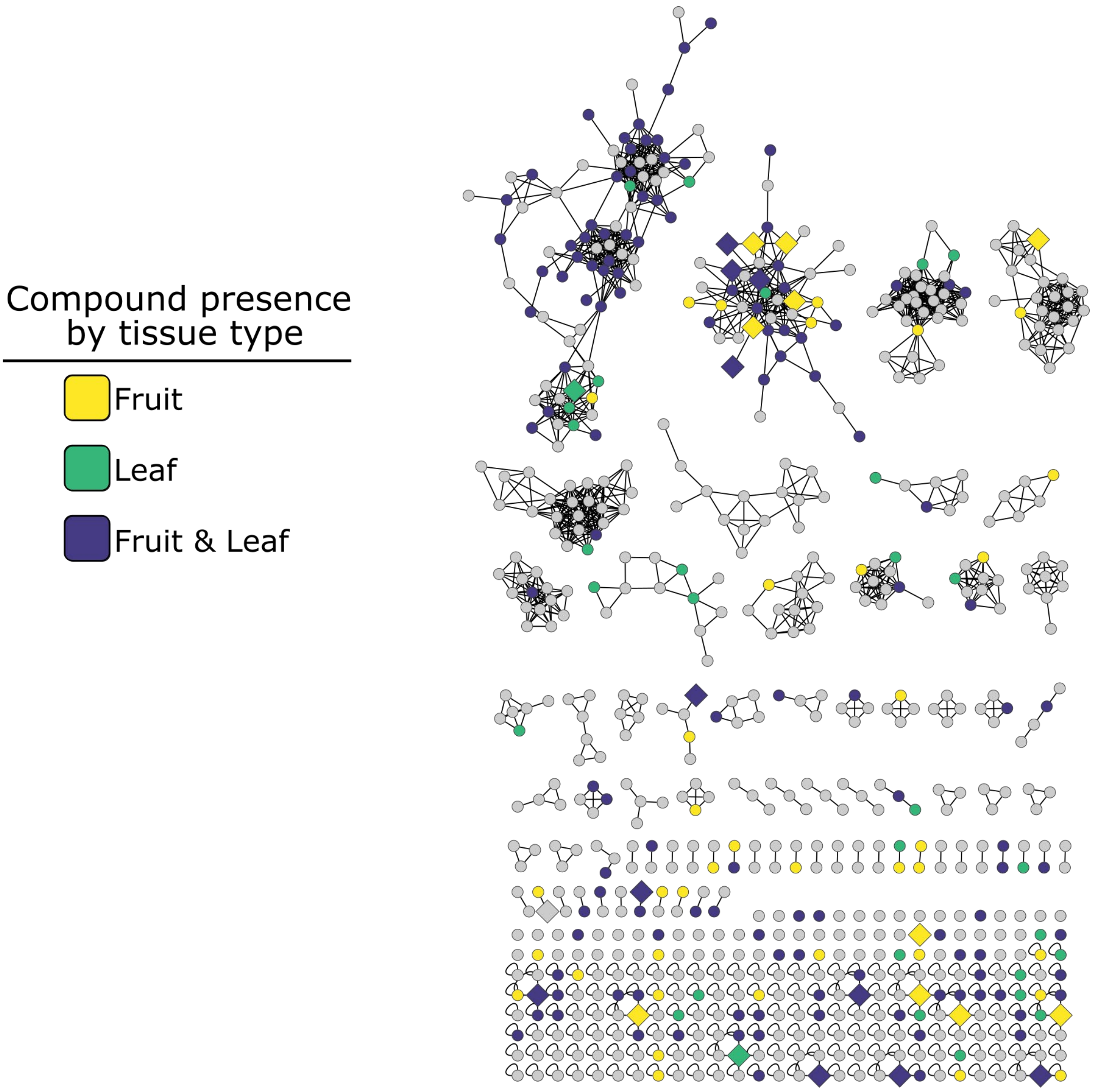


34

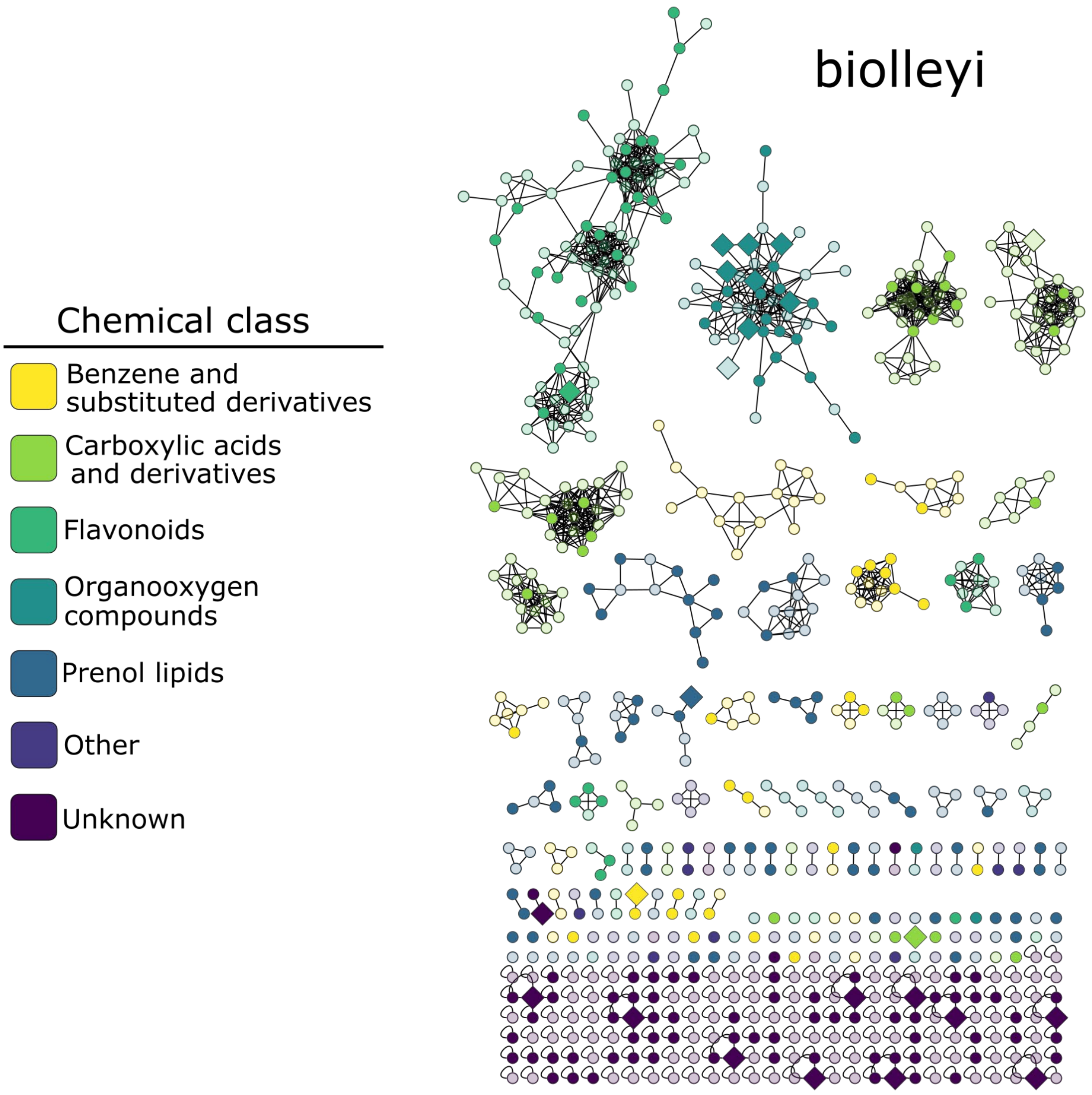
Chemical class
Benzene and substituted derivatives
Carboxylic acids and derivatives
Flavonoids
Organooxygen compounds
Prenol lipids
Other
Unknown
biolleyi

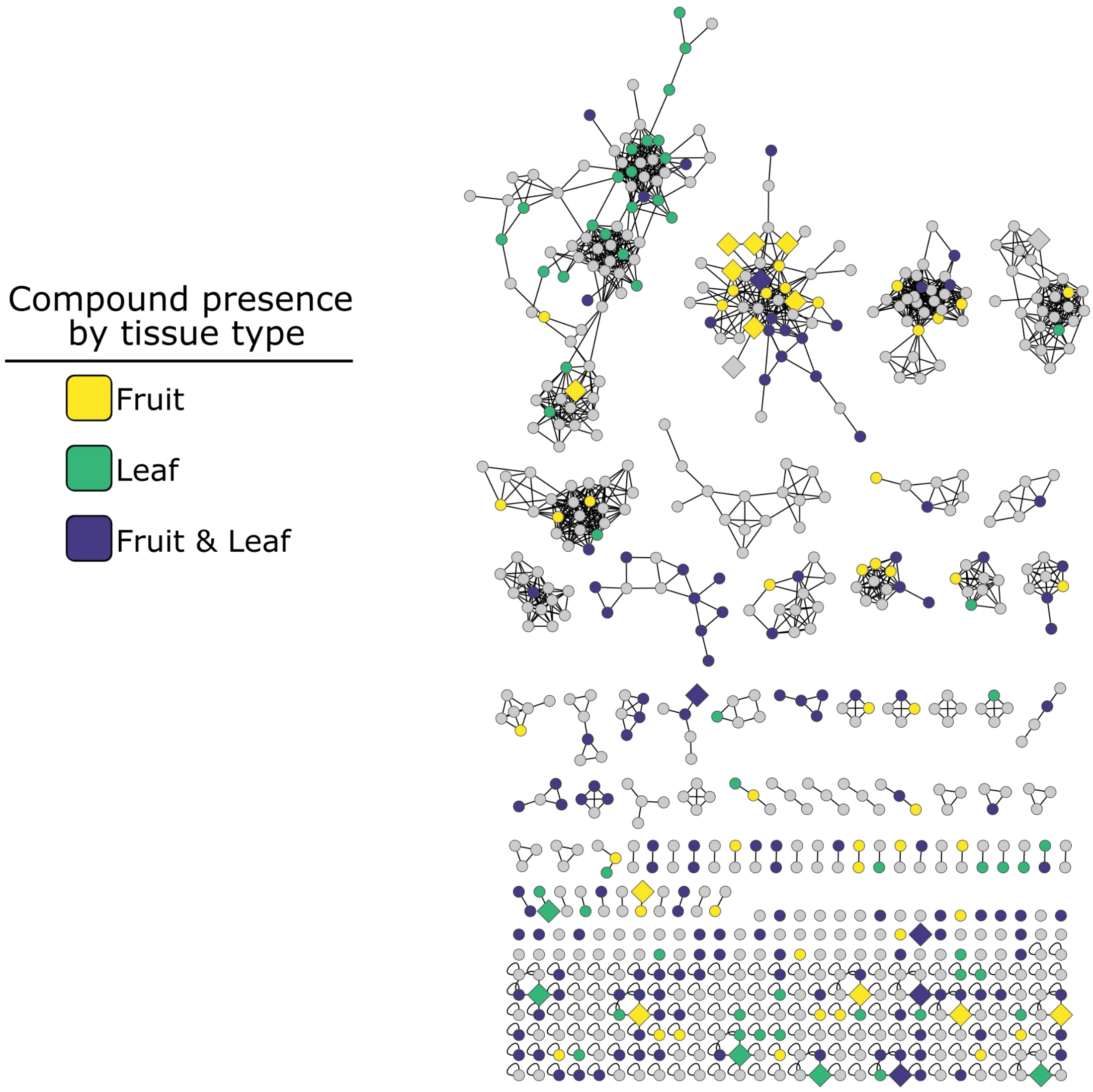


37

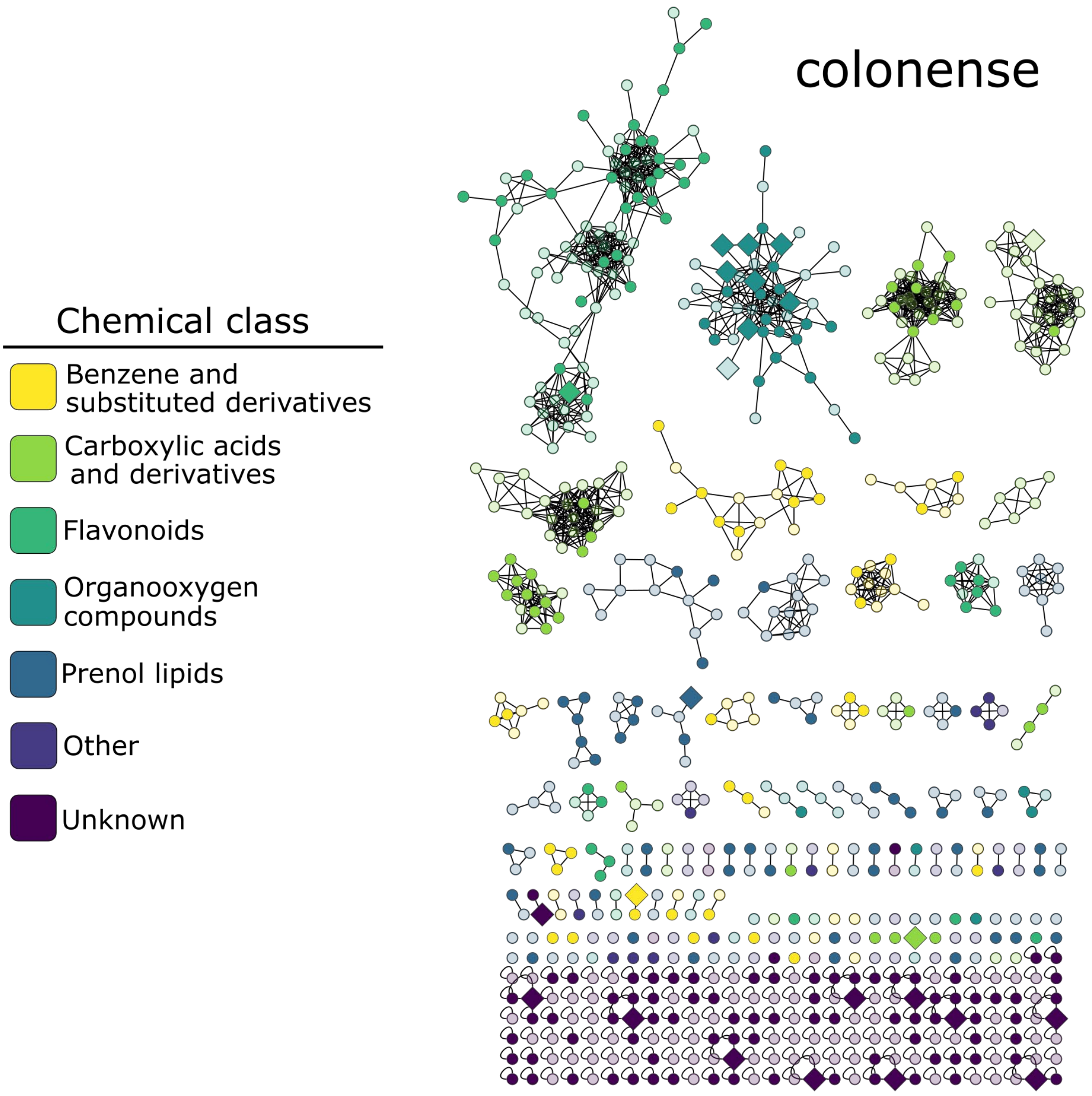
colonense
Chemical class
Benzene and substituted derivatives
Carboxylic acids and derivatives
Flavonoids
Organooxygen compounds
Prenol lipids
Other
Unknown

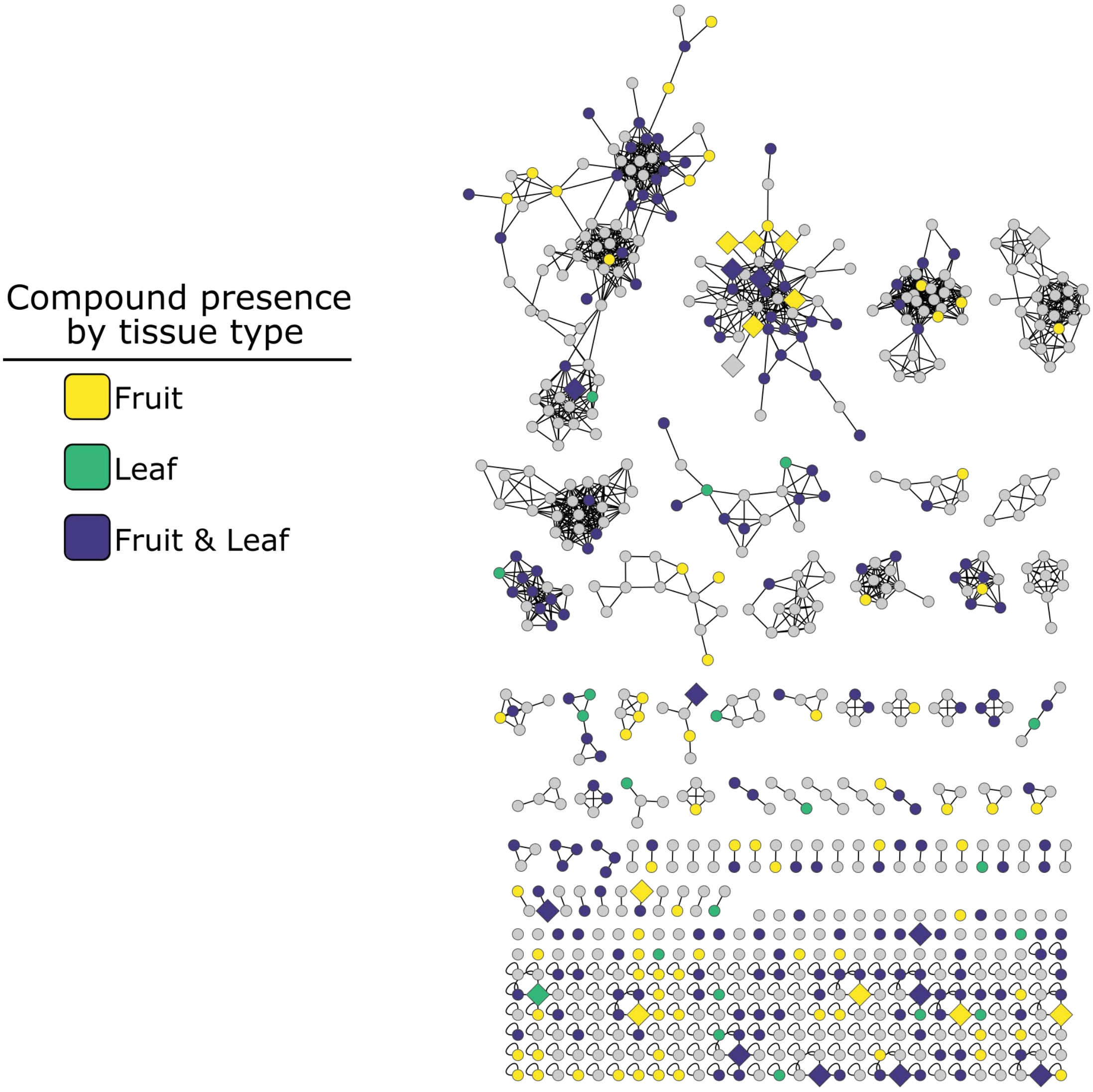
Compound presence by tissue type
Fruit
Leaf
Fruit & Leaf

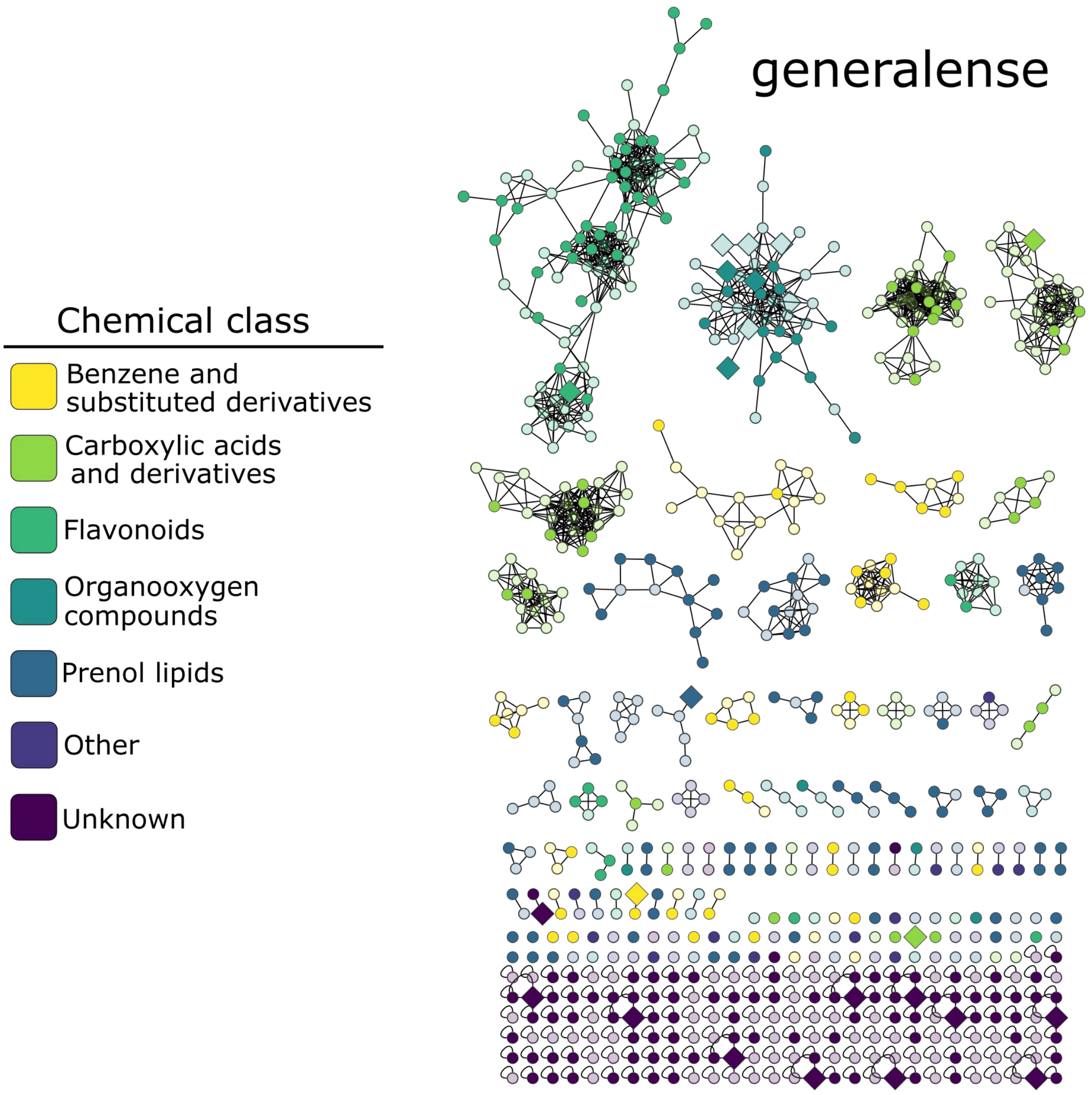
generalense
Chemical class
Benzene and substituted derivatives
Carboxylic acids and derivatives
Flavonoids
Organooxygen compounds
Prenol lipids
Other
Unknown

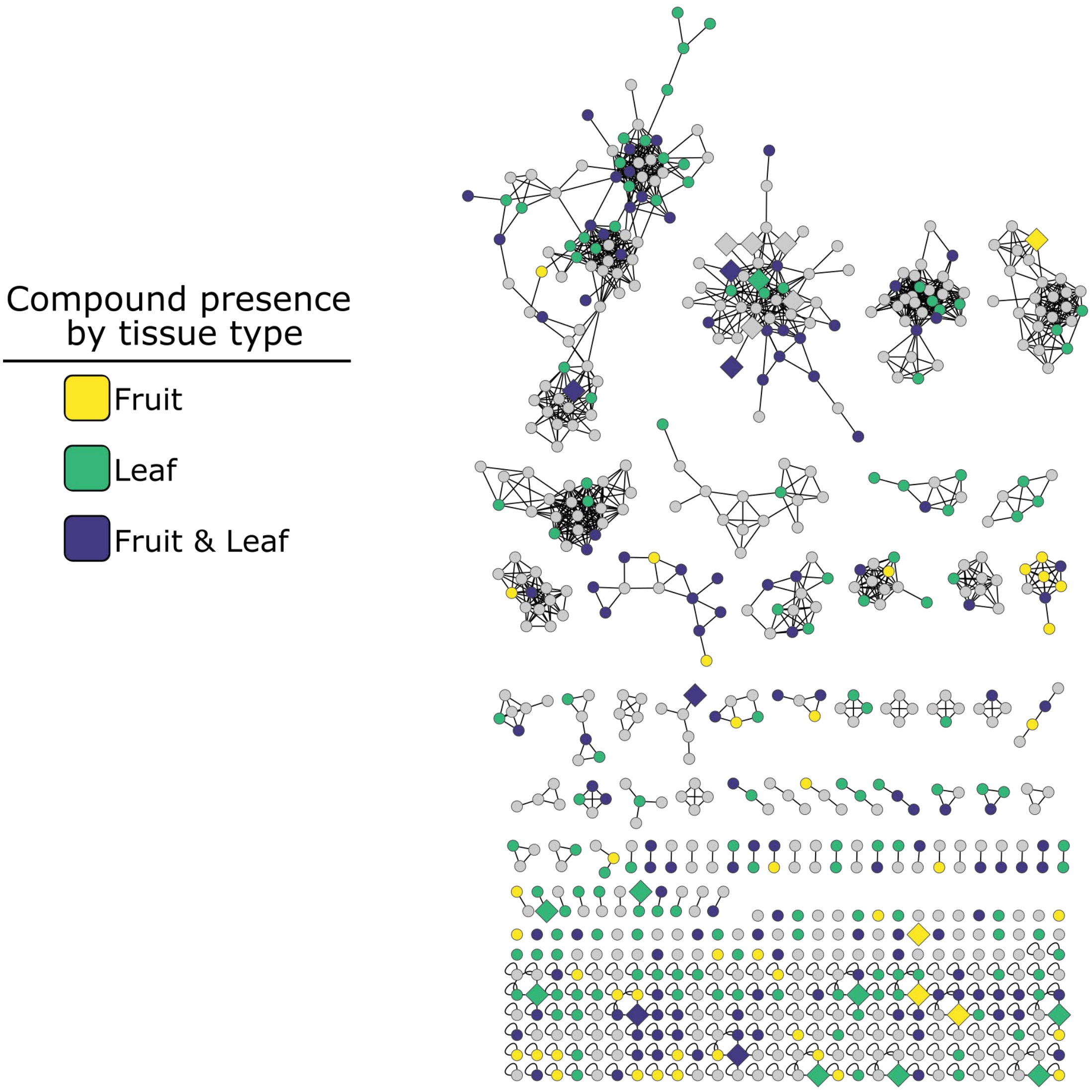
Compound presence by tissue type
Fruit
Leaf
Fruit & Leaf

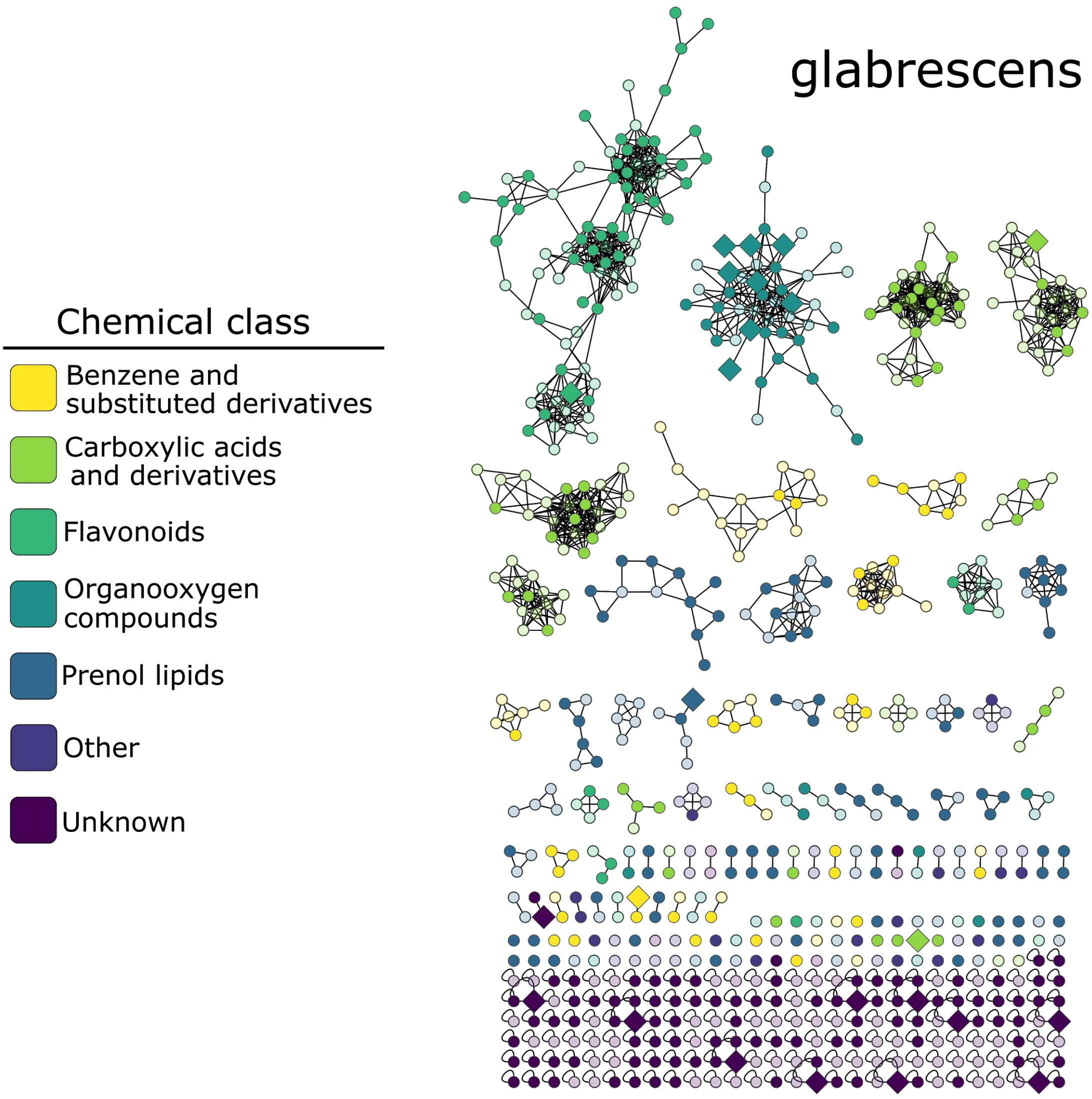
glabrescens
Chemical class
Benzene and substituted derivatives
Carboxylic acids and derivatives
Flavonoids
Organooxygen compounds
Prenol lipids
Other
Unknown

50

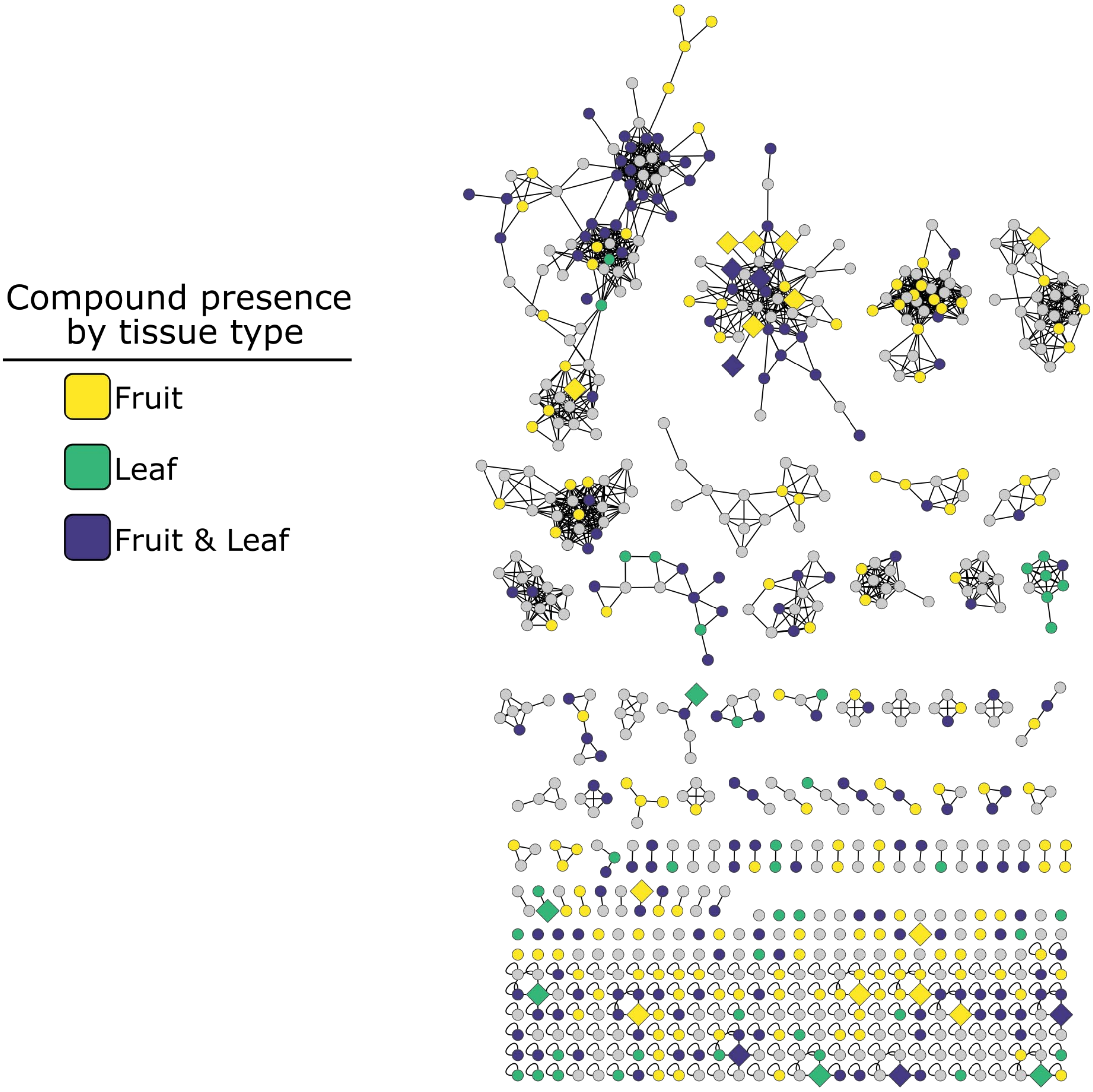


51

52

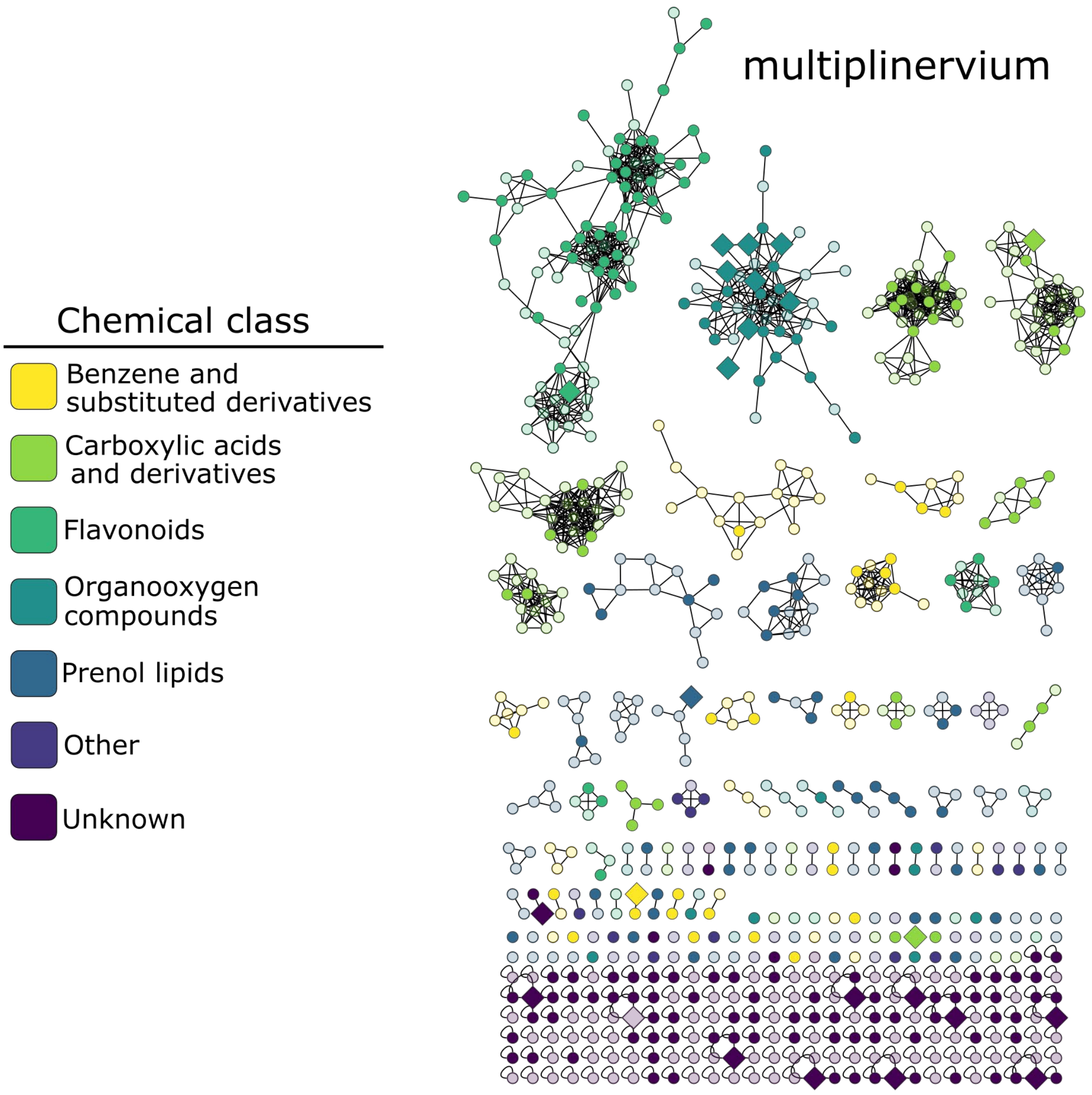
Chemical class
Benzene and substituted derivatives
Carboxylic acids and derivatives
Flavonoids
Organooxygen compounds
Prenol lipids
Other
Unknown
multiplinervium

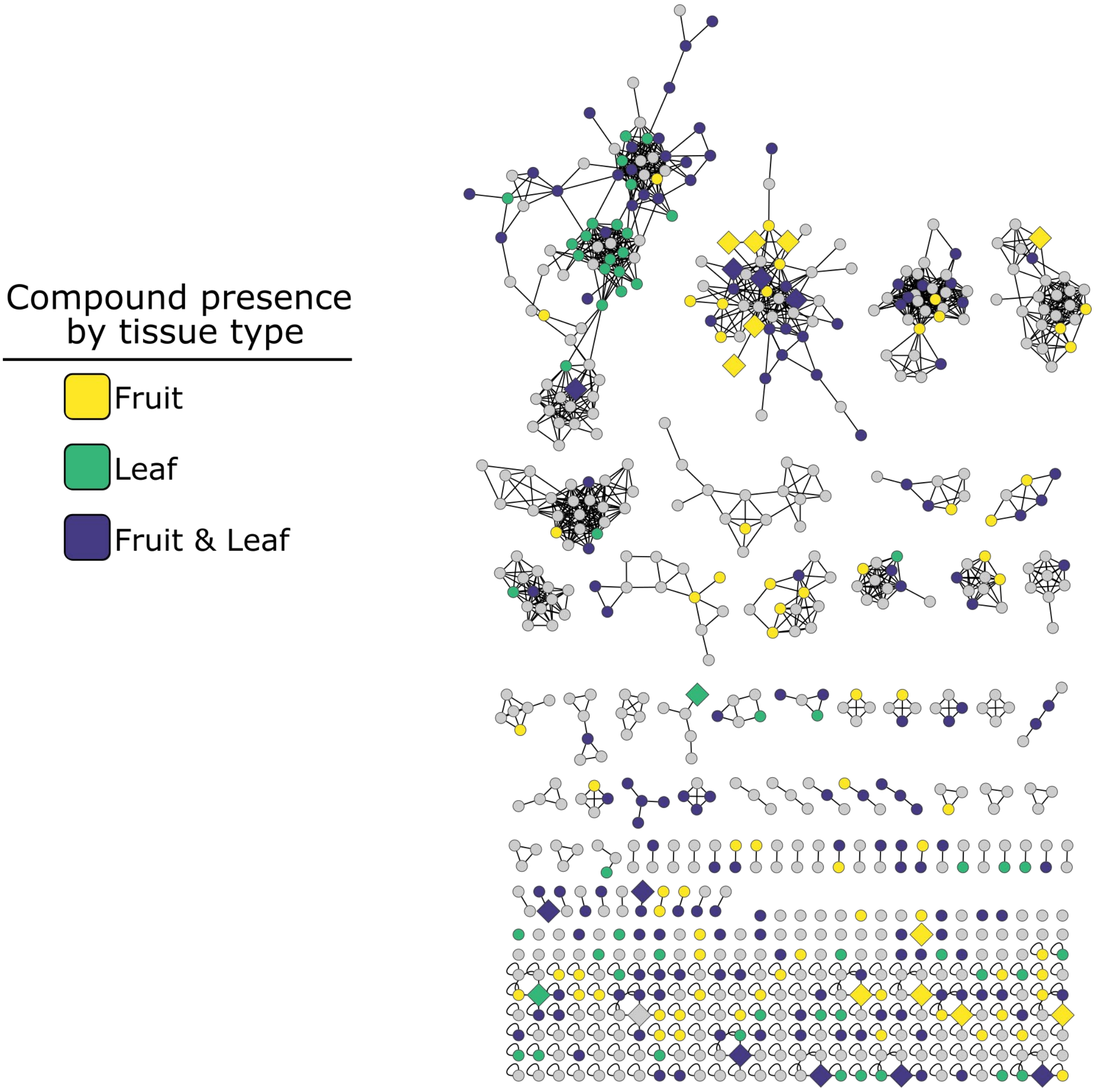
Compound presence
by tissue type
Fruit
Leaf
Fruit & Leaf

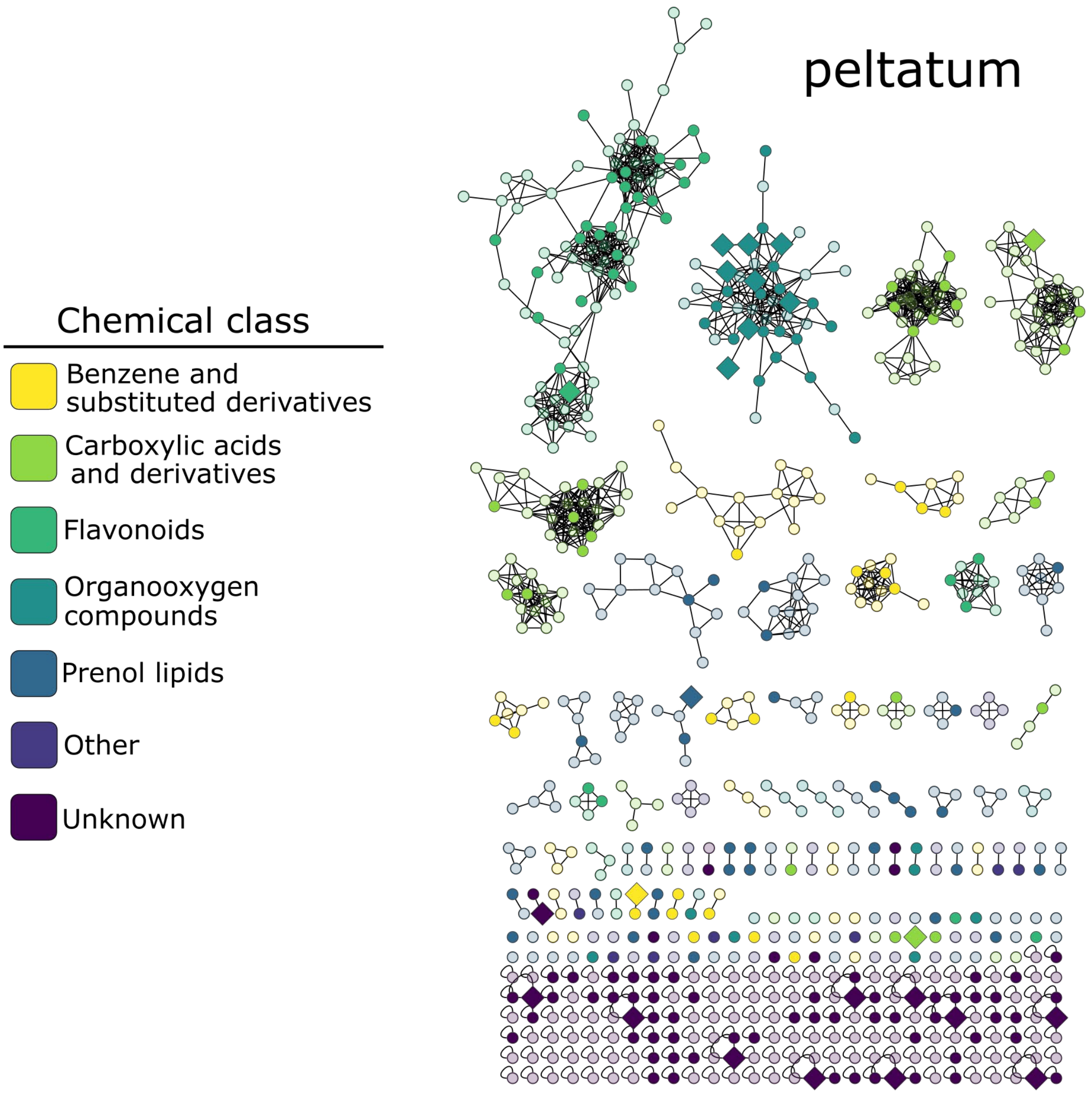
peltatum
Chemical class
Benzene and substituted derivatives
Carboxylic acids and derivatives
Flavonoids
Organooxygen compounds
Prenol lipids
Other
Unknown

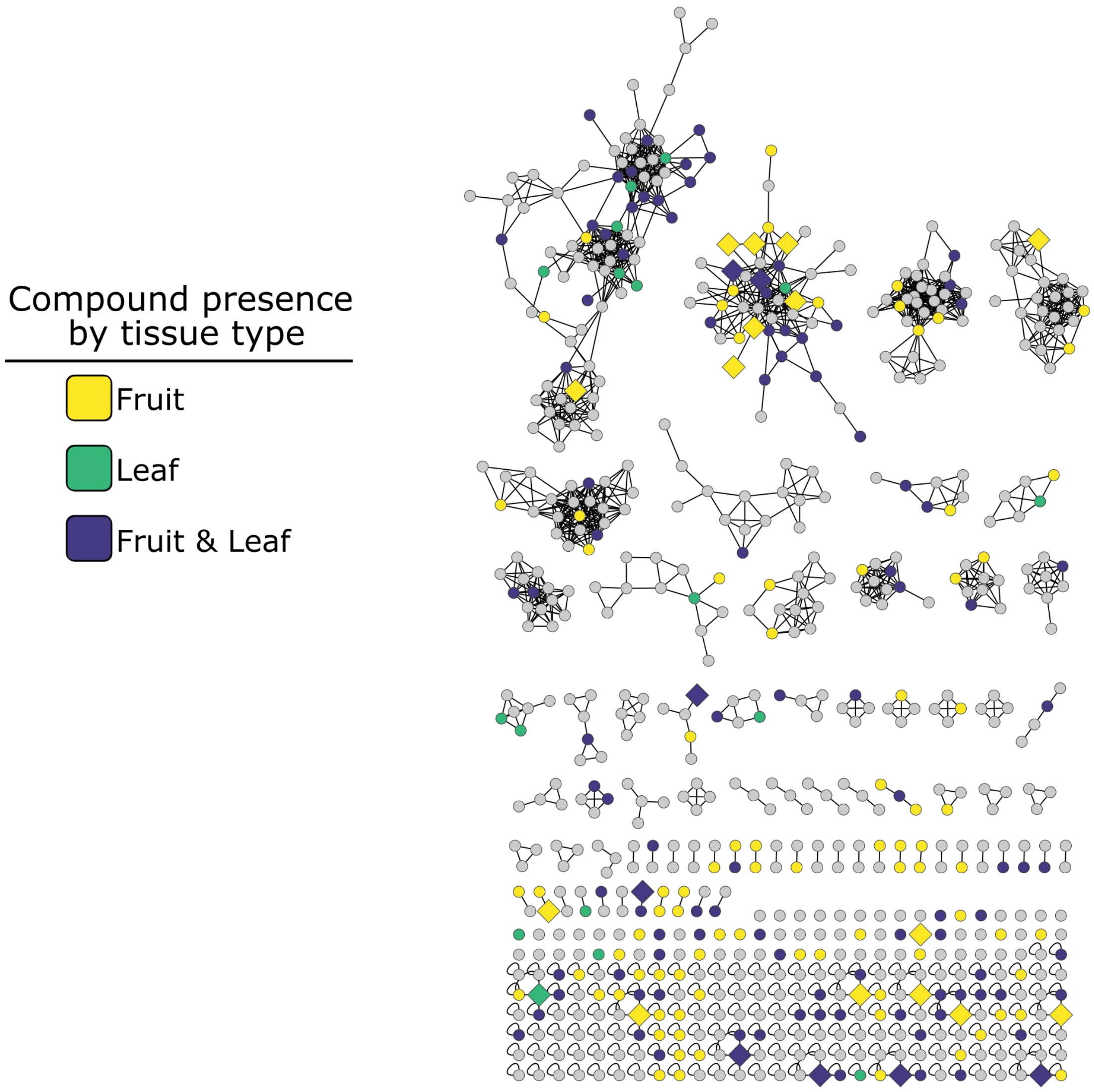
Compound presence
by tissue type
Fruit
Leaf
Fruit & Leaf

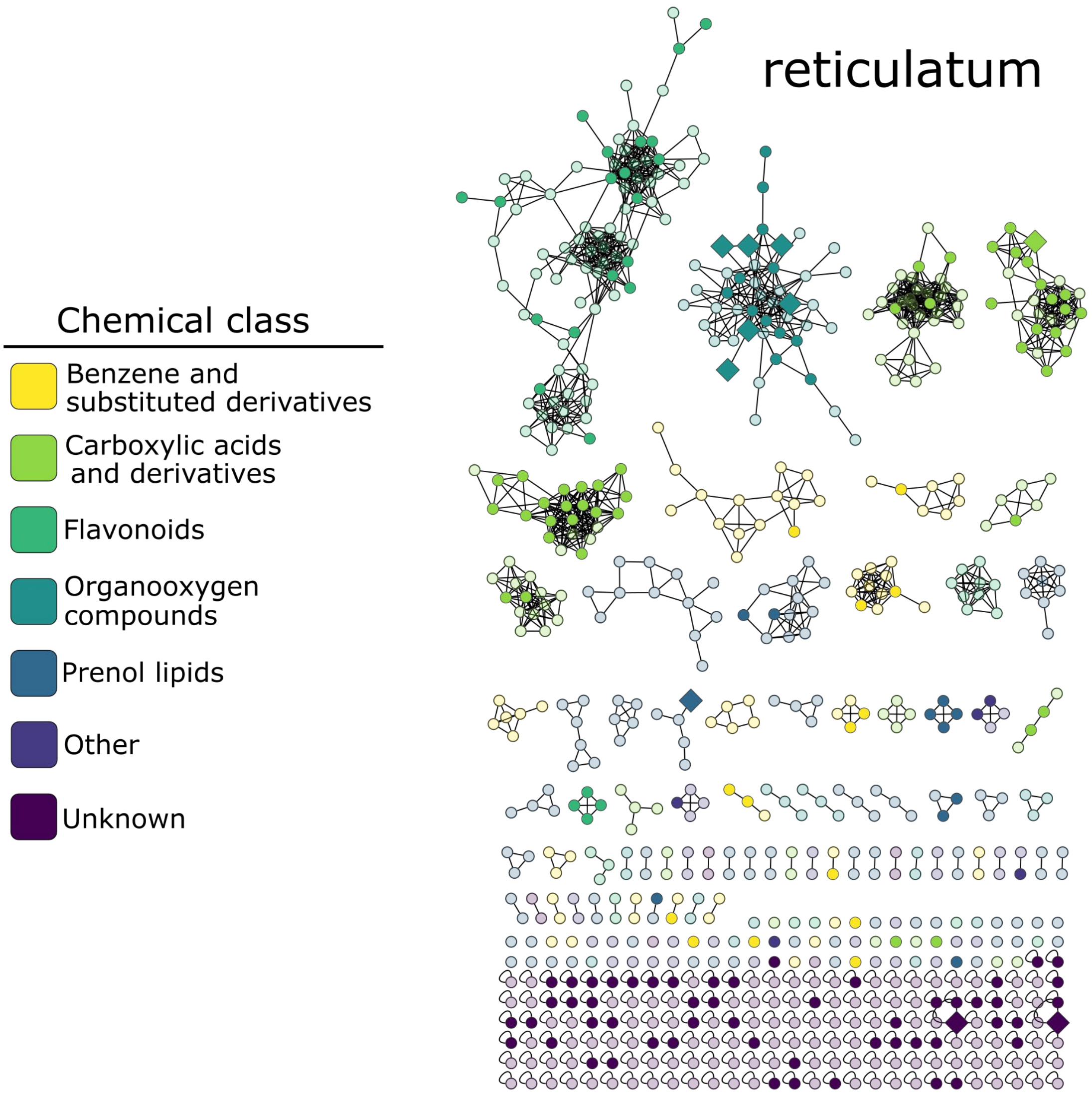
reticulatum
Chemical class
Benzene and substituted derivatives
Carboxylic acids and derivatives
Flavonoids
Organooxygen compounds
Prenol lipids
Other
Unknown

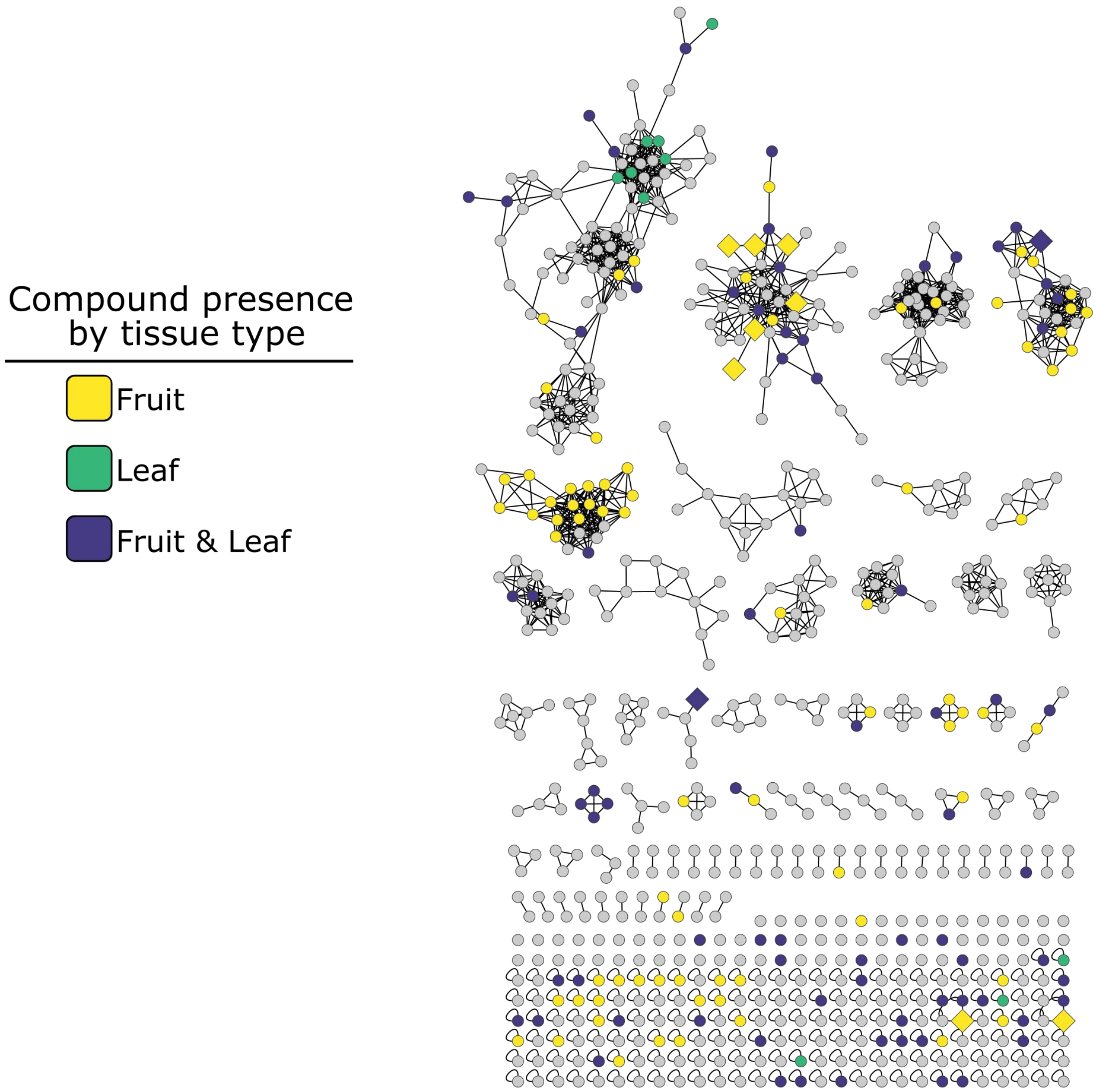

Compound presence by tissue type
Fruit
Leaf
Fruit & Leaf

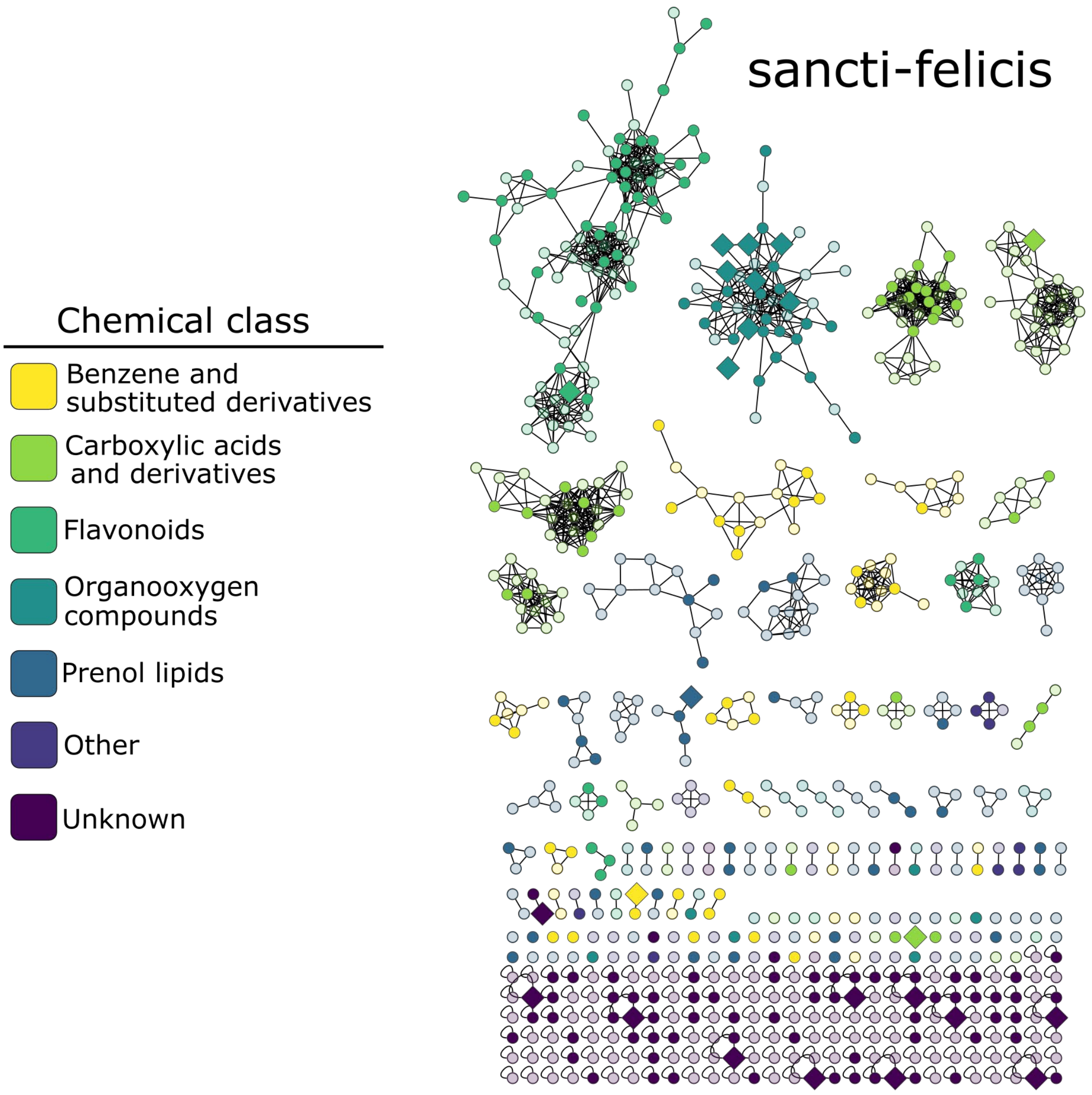
sancti-felicis
Chemical class
Benzene and substituted derivatives
Carboxylic acids and derivatives
Flavonoids
Organooxygen compounds
Prenol lipids
Other
Unknown

62

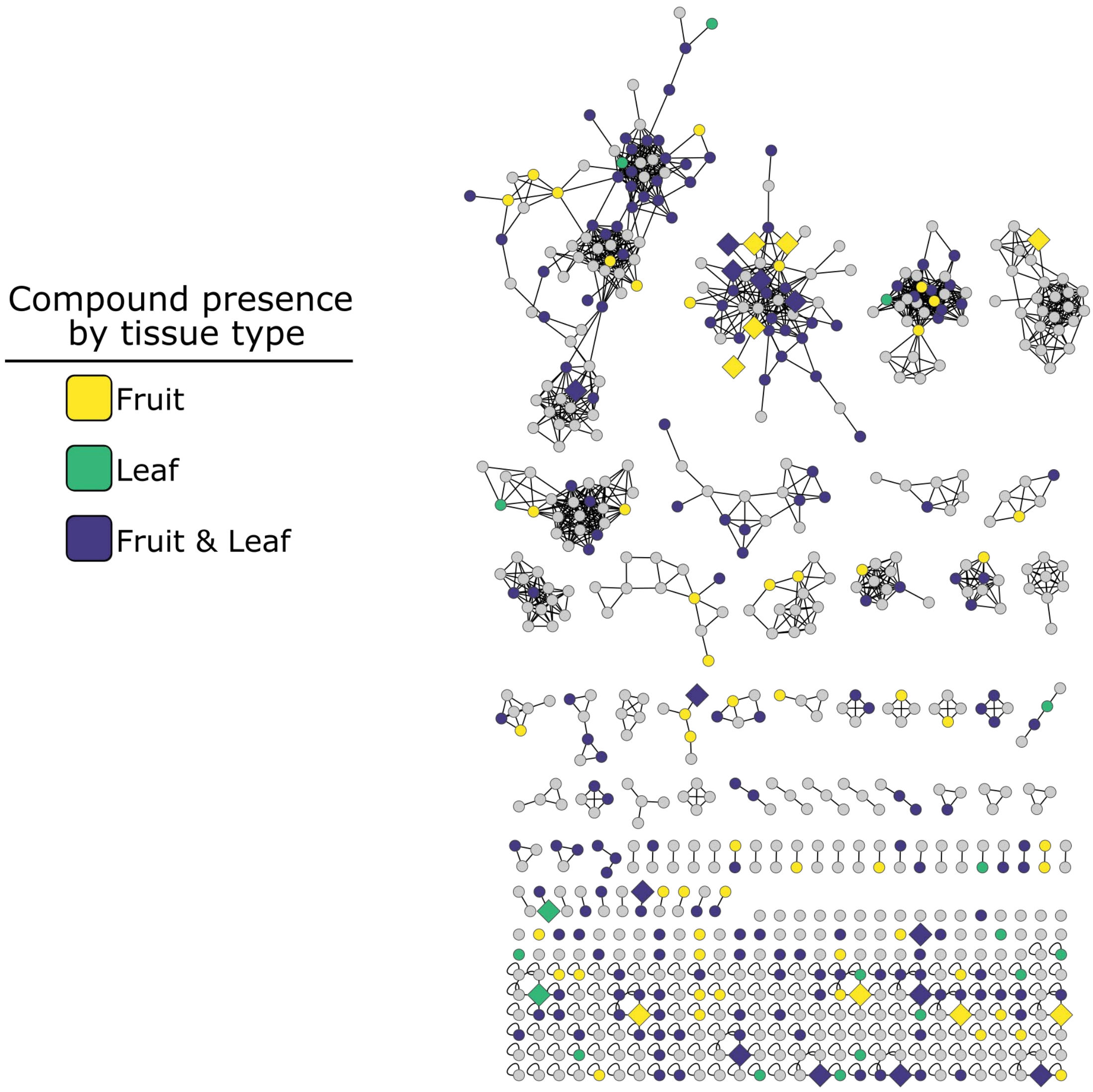
Compound presence
by tissue type
Fruit
Leaf
Fruit & Leaf

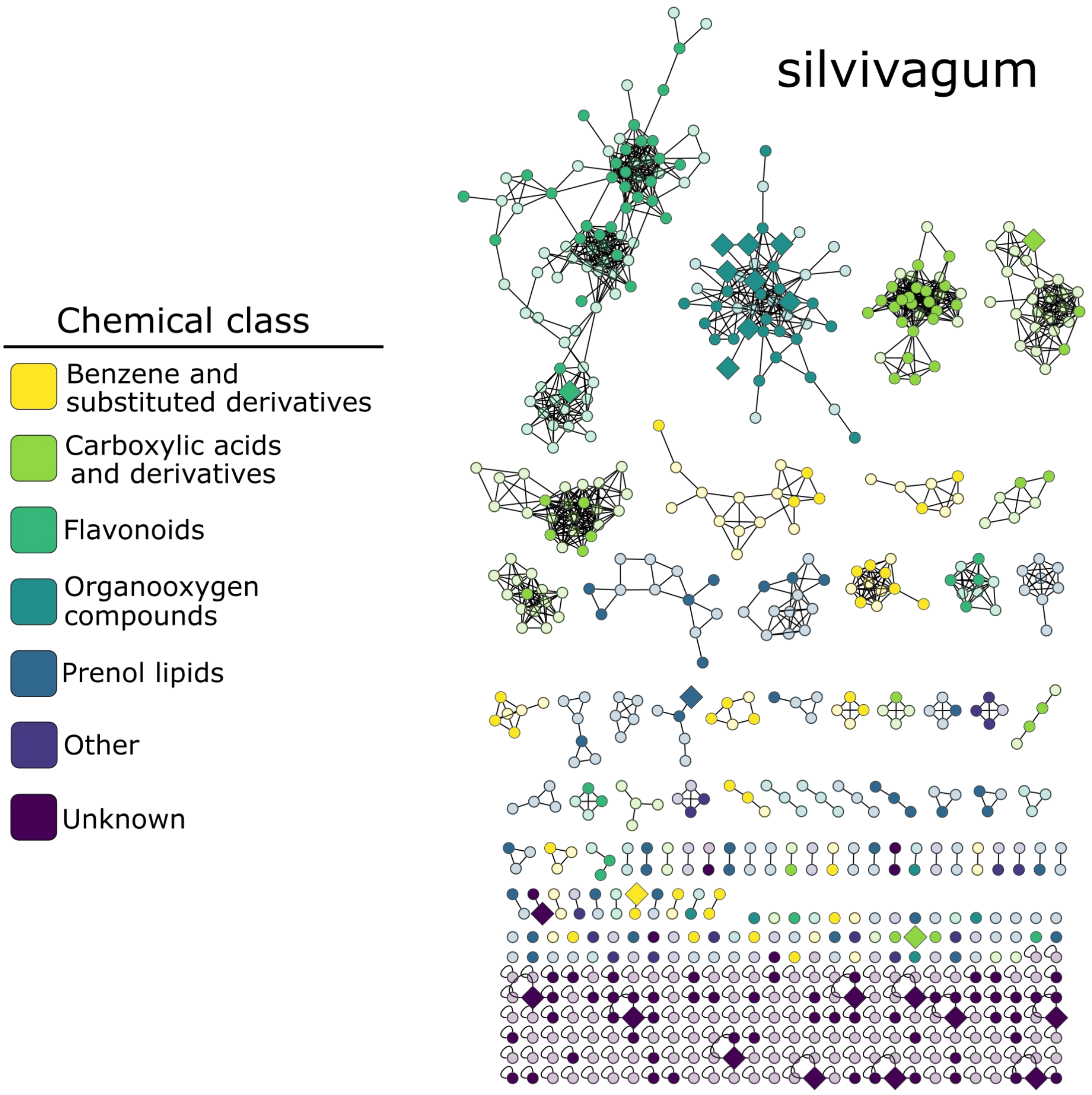
silvivagum
Chemical class
Benzene and substituted derivatives
Carboxylic acids and derivatives
Flavonoids
Organooxygen compounds
Prenol lipids
Other
Unknown

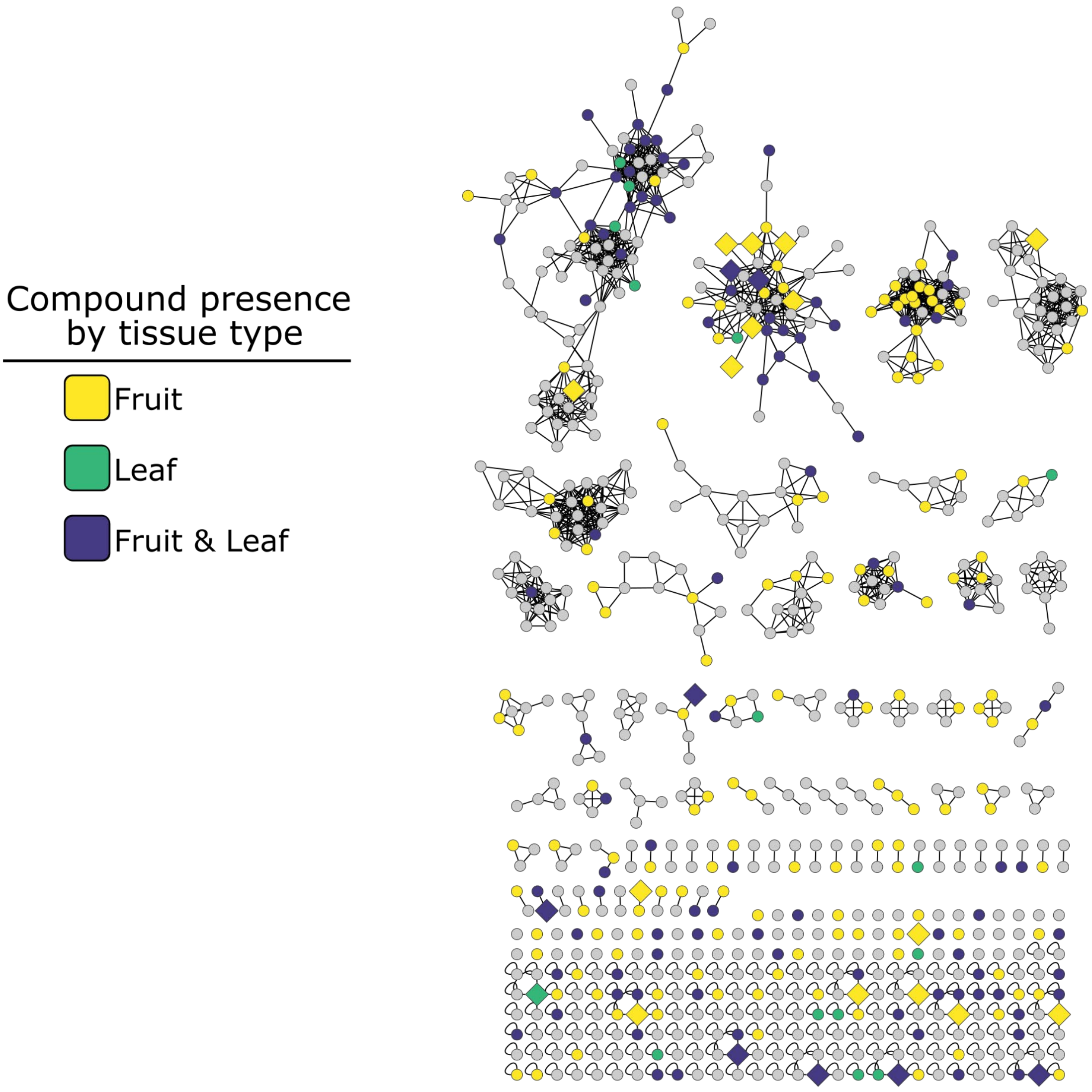
Compound presence
by tissue type
Fruit
Leaf
Fruit & Leaf

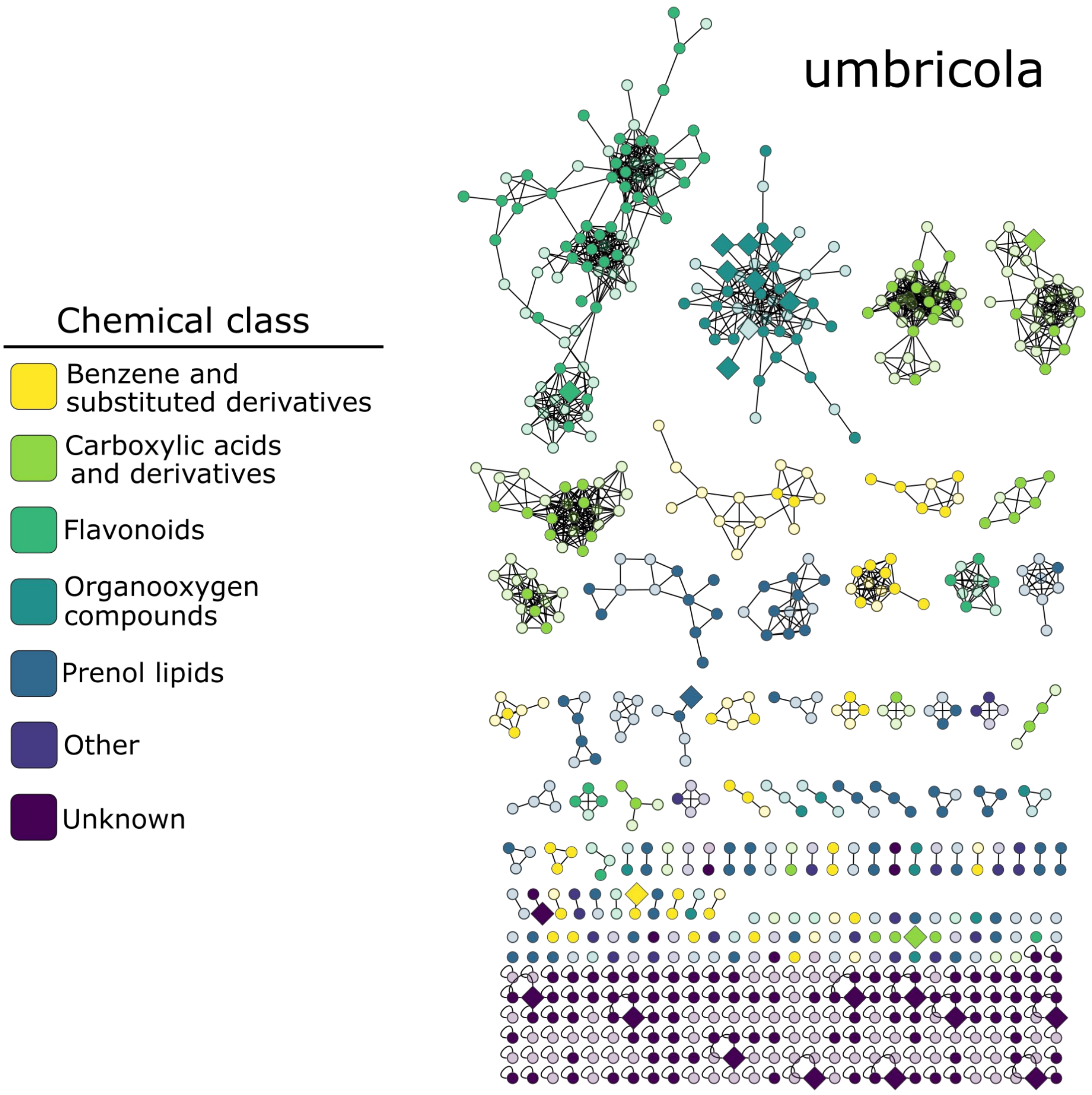
umbricola
Chemical class
Benzene and substituted derivatives
Carboxylic acids and derivatives
Flavonoids
Organooxygen compounds
Prenol lipids
Other
Unknown

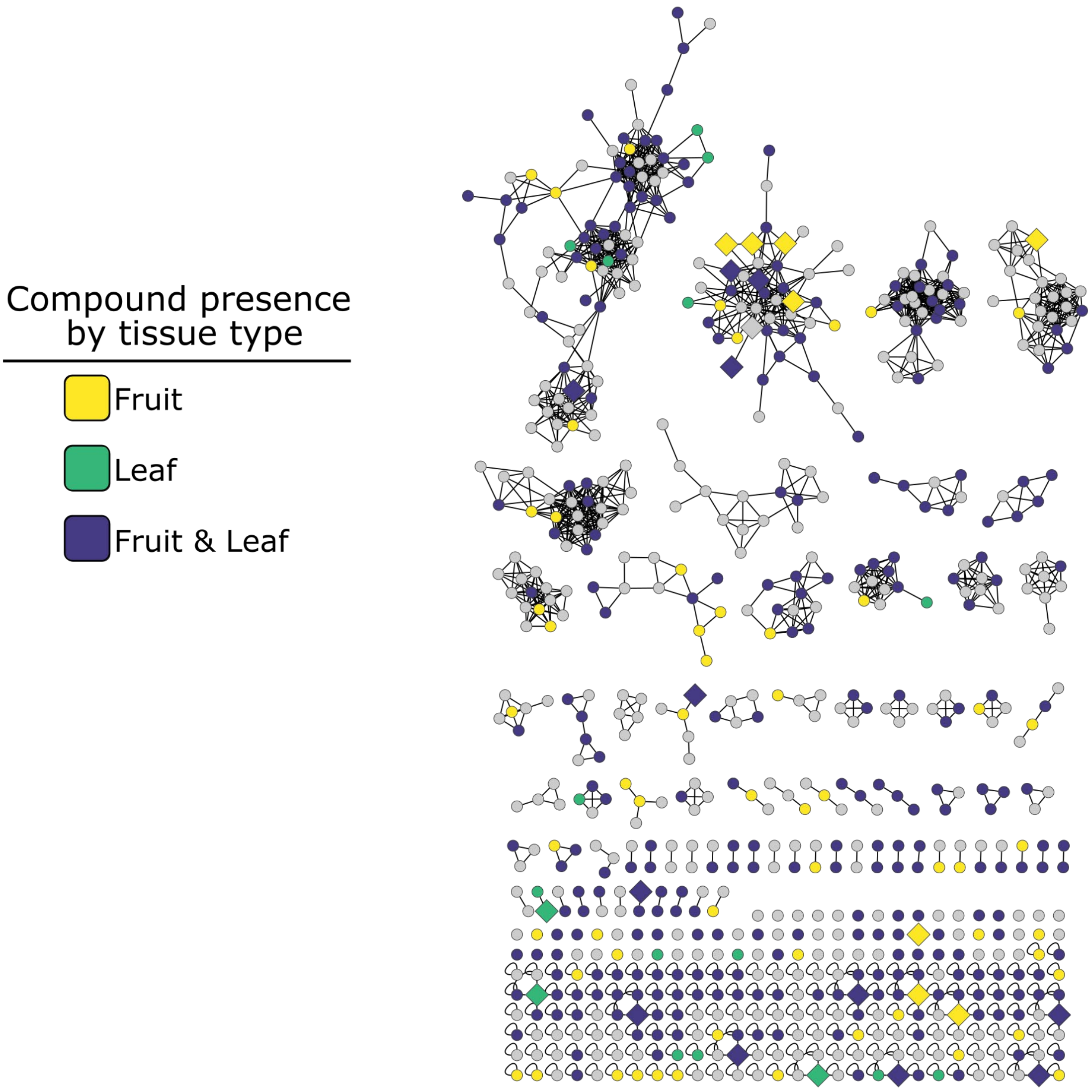
Compound presence
by tissue type
Fruit
Leaf
Fruit & Leaf
